# Dynamic allostery drives acetyl-CoA-mediated activation of *Mycobacterium tuberculosis* isocitrate lyase 2

**DOI:** 10.1101/2025.08.02.651367

**Authors:** Evelyn Yu-Wen Huang, Brooke X.C. Kwai, Wanting Jiao, Jamie Taka, Karyn L. Wilde, Ashish Sethi, Megan J. Maher, Ghader Bashiri, Ivanhoe K.H. Leung

**Author notes:** To whom correspondence should be addressed. Email addresses. These authors contributed equally to this paper (Evelyn Yu-Wen Huang, Brooke X.C. Kwai). **Author Contributions:** E.Y.W.H., B.X.C.K., and I.K.H.L. designed research. E.Y.W.H., B.X.C.K., W.J., and M.J.M. performed research. E.Y.W.H., B.X.C.K., W.J., A.S., and M.J.M. analysed data. K.L.W., A.S., and M.J.M. contributed new tools/techniques. E.Y.W.H., B.X.C.K., W.J., G.B., J.T., M.J.M. and I.K.H.L. wrote the paper. **Data Deposition:** The atomic coordinates and structure factors have been deposited in the Protein Data Bank, www.pdb.org (PDB ID 9OBO). The chemical shift assignments for the ICL2 C-terminal domain (ICL2_601-766_) have been deposited in the BioMagResBank (http://bmrb.io) under the accession numbers 52666 and 52668.

## Abstract

*Mycobacterium tuberculosis* isocitrate lyase 2 (ICL2) is an allosterically regulated enzyme that enables the bacterium to survive on non-glycolytic substrates during infection. Previous studies showed that ICL2 is allosterically regulated by acetyl-CoA and its analogues but the molecular mechanism underpinning this regulation is unknown. Here, we use protein NMR, crystallography, molecular dynamics, and mutagenesis studies to show that two unique structural features of ICL2, its C-terminal domain and a unique helical substructure on its N-terminal catalytic domain, play important roles in the enzyme’s allostery. In particular, we found that the binding of acetyl-CoA promotes the dimerisation of the C-terminal domain and disrupts its interactions with the unique helical substructure on the N-terminal domain. This leads to conformational changes in the ICL2 enzyme that induces activation. Taken together, our findings reveal, for the first time, how the binding of acetyl-CoA, which is not an ICL2 substrate, induces ICL2 activation. By extension, the work also identifies a novel allosteric mechanism controlling *M. tuberculosis* metabolism that is amenable to therapeutic manipulation.

**Significance Statement:** *Mycobacterium tuberculosis* isocitrate lyase 2 (ICL2) was previously shown to be activated by acetyl-CoA and propionyl-CoA – two central metabolites generated by the metabolism of sugars and fatty acids. However, it is not known how the binding of these metabolites leads to the activation of ICL2. Together with its isoform ICL1, ICL2 has been shown to be essential for the survival and pathogenesis of the bacterium. Understanding how this regulation occurs can help design novel treatments to target this protein and eradicate these bacteria, which cause the most deaths worldwide due to a single bacterial agent. This system also presents a fascinating model to examine allostery in proteins, with the techniques illustrated in this paper being applicable to other allosteric proteins.

## Introduction

The ability of cells to modulate the activity and function of their enzymes is crucial for maintaining homeostasis and adapting to environmental changes. This regulation is achieved through multiple mechanisms, including, but not limited to, gene expression, post-translational modifications, allosteric regulation, and inhibition. Among these, allostery, which is the perturbation of a protein at one site that leads to an effect at another site, allows rapid, reversible control of enzyme function in response to metabolic signals, such as changes in the concentration of small molecule metabolites or biomolecules [1-5]. Understanding the mechanism behind allostery provides fundamental insights into how proteins coordinate internal communication to regulate physiological processes. It could also inform the development of allosteric modulators for applications such as drug discovery, a relatively less explored but growing approach for the development of new therapeutics [6-8].

A notable example of an allosteric enzyme is isocitrate lyase (ICL) isoform 2 of *Mycobacterium tuberculosis* (*Mtb*). ICL is the first enzyme of the glyoxylate shunt that catalyses the aldol cleavage of isocitrate to form succinate and glyoxylate (Fig. 1a). Its activity is crucial for the ability of the bacterium to survive on non-glycolytic carbon sources (such as fatty acids and acetate) during infection. In *Mtb*, there are two isoforms of ICL (ICL1 and ICL2) [9]. The activity of *Mtb* ICL2 is allosterically regulated by acetyl-CoA, propionyl-CoA and other CoA analogues [10], which are key metabolites that are produced during fatty acid metabolism. Inhibition of ICL has been proposed as a strategy to treat tuberculosis, as knockout studies have revealed these enzymes to be essential for bacterial survival and virulence [11]. However, due to the small and polar nature of the active sites of these enzymes, no ICL inhibitors have progressed beyond the initial preclinical development stage [12]. Targeting ICL2 allostery therefore represents an alternative strategy to modulate activity.

**Figure 1.**
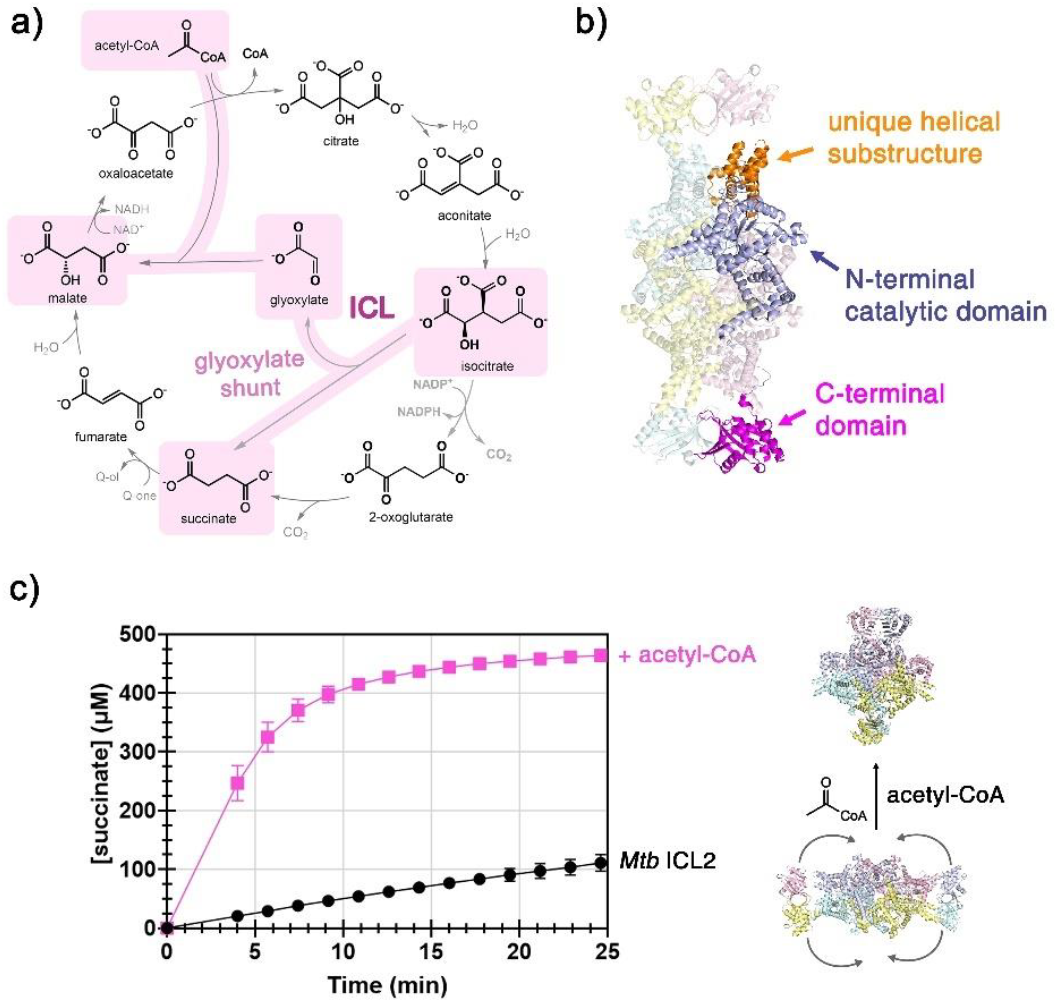
Unique structure and activation of *M. tuberculosis* (*Mtb*) ICL2. (a) The glyoxylate shunt within the tricarboxylic acid (TCA) cycle. The glyoxylate shunt bypasses the two decarboxylation steps in the TCA cycle. ‘ICL’ indicates the ICL-catalysed reaction in the glyoxylate shunt. (b) Structure of *Mtb* ICL2 (PDB: 6EDW [10]) which forms a homotetramer. Each monomer is indicated by a different colour with one monomer coloured according to the three features present in each monomer. (c) The acetyl-CoA-mediated activation of ICL2. ICL2 is activated by acetyl-CoA, and this activation is accompanied by conformational changes in the ICL2 structure. The *Mtb* ICL2 structure in the absence and presence of acetyl-CoA was taken from PDB 6EDW [10] and 6EE1 [10], respectively.

Structurally, *Mtb* ICL2 is distinct from ICLs found in most other organisms. Based on bioinformatic analyses, ICLs are classified into three structural classes (Fig. S1) [11]. Group I ICLs, such as *Mtb* ICL1, contain a single catalytic domain with the conserved KKCGH motif that is required for their catalytic activity. Group II ICLs are also single-domain enzymes with the same quaternary structure as Group I ICLs. However, their catalytic domain contains an additional helical substructure that extends the domain. These ICLs are exclusively found in eukaryotes such as plants and fungi [11], and it is thought that the helical substructure is responsible for cellular localisation [13]. Group III ICLs are multidomain enzymes that so far have only been observed in mycobacteria. *Mtb* ICL2 is the only well-characterised member in this group [10].

*Mtb* ICL2 forms a homo-tetramer (Fig. 1b), with each subunit contains an N-terminal catalytic domain (residues 1-582) that comes together to form the core of the tetramer. Each catalytic domain also carries a helical substructure (residues 278–428) similar to Group II ICLs. However, the function of this helical substructure is not known. It is unlikely that this substructure would be responsible for localisation since *Mtb* lacks compartmentalisation. Extending from the catalytic domain is a unique C-terminal domain (CTD, residues 601–766) which is not found in canonical ICLs and is connected to the catalytic core via a flexible linker (residues 582–600). Binding of acetyl-CoA triggers conformational changes that involve the reorganisation of the CTDs, resulting in a 50-fold increase in catalytic efficiency (Fig. 1c) [10]. However, the mechanism by which the binding of acetyl-CoA induces conformational changes and activates the enzyme remains unknown.

Herein, we report how the unique structural features in *Mtb* ICL2 contribute to its function and how different CoA analogues modify these structural features to activate ICL2. We found that the unique helical substructure and the CTD work together to mediate the acetyl-CoA-mediated activation of *Mtb* ICL2. Our results therefore provide the first mechanistic insight into this novel allosteric regulatory process in *Mtb* and also identify a new allosteric target for anti-tuberculosis therapies.

## Results

### The C-terminal domain of Mtb ICL2 is required for acetyl-CoA-mediated activation

We first investigated the role of the CTD in the catalytic activity and allostery of ICL2 by generating a truncated variant ICL2_1-600_ (ICL2ΔCTD) which lacks the CTD. A previous study also used a similar construct of *Mtb* ICL2 (ICL2_1-605_) to test the efficacy of a covalent-based inhibitor [14]. In the presence of Mg^2+^ (a co-factor required for all ICL reactions), ICL2ΔCTD was active with a reaction rate that is similar to that of the full length ICL2 under the same conditions. However, while full length ICL2 is regulated allosterically, the activity of ICL2ΔCTD was not accelerated by acetyl-CoA (Fig. 2a). These results show that the basal activity of ICL2 has no correlation with the CTD, but the CTD is necessary for the activation of ICL2 mediated by acetyl-CoA.

**Figure 2.**
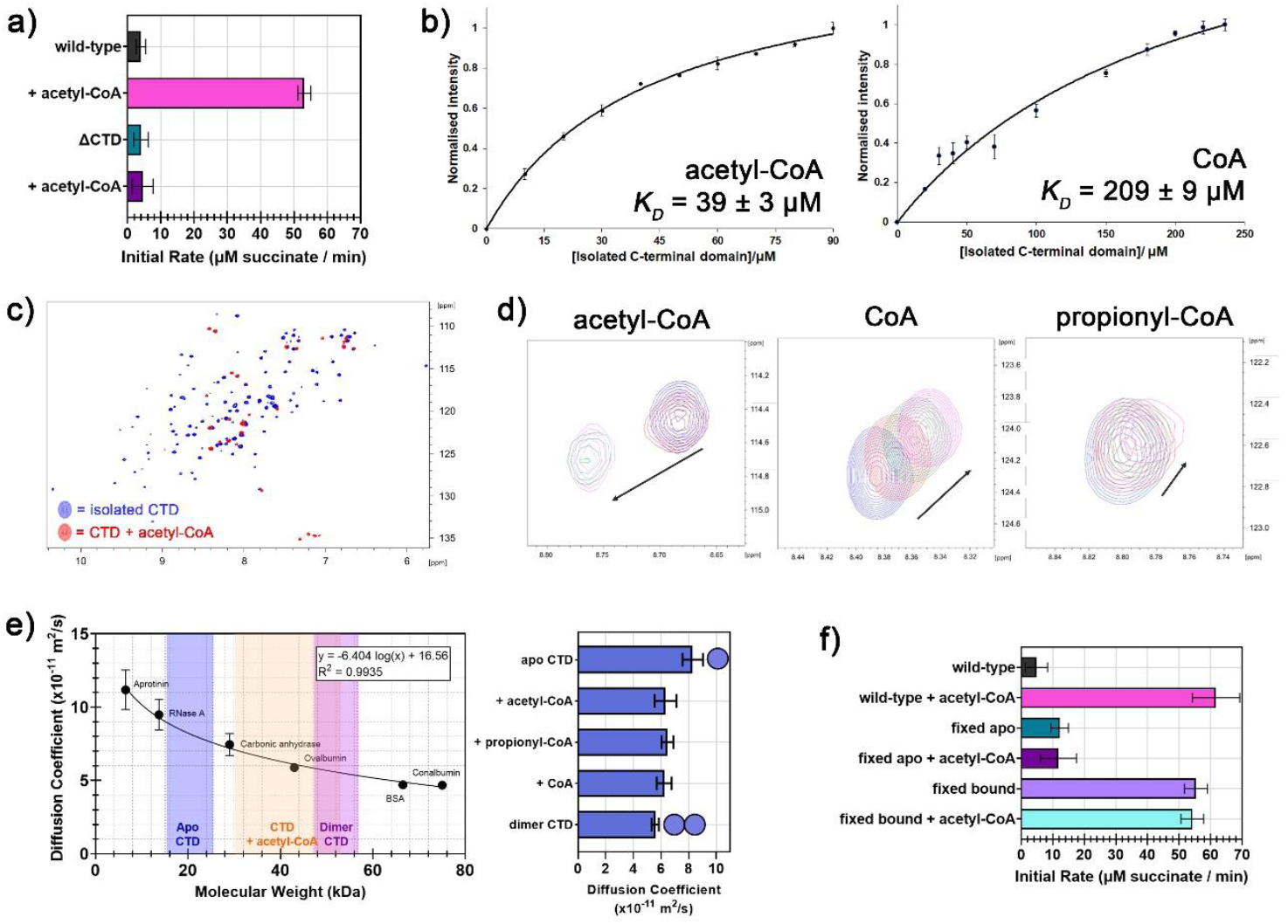
The role of the C-terminal domain (CTD) in *M. tuberculosis* ICL2. Enzyme kinetics measurements were performed at 298 K, and samples contained 500 nM enzyme, 1 mM DL-isocitrate, 5 mM MgCl2, 25 μM acetyl-CoA (where applicable), 50 mM Tris-d11 (pH 7.5), 0.02% NaN3 in 90% H2O / 10% D_2_O. (a) Effect of CTD deletion on ICL2 activity. Error bars represent standard deviation from three measurements. (b) Binding curves between ICL2 CTD (variable concentration) and acetyl-CoA (50 μM) or CoA (50 μM). Experiments were conducted in 50 mM Tris-d11 (pH 7.5) in 90% H_2_O / 10% D_2_O at 298 K. Error bars represent standard deviation from three separate experiments. (c) Effect of acetyl-CoA binding on the HSQC spectrum of ^15^N-labelled ICL2 CTD. Overlay of ^1^H-^15^N HSQC spectra of 40 µM^15^N-ICL2 CTD in the absence (blue) and presence (red) of 2 mM acetyl-CoA. (d) Titration of acetyl-CoA, CoA, or propionyl-CoA to ICL2 CTD (^2^H, ^15^N-labelled). For acetyl-CoA, a zoomed region on residue S650 of overlaid ^1^H-^15^N HSQC spectra for the 80 µM CTD titrated with acetyl-CoA at a concentration of 0 µM (blue), 50 µM (red), 100 µM (green) and 175 µM (magenta) with an arrow indicating the direction of chemical shift perturbation during titration. For CoA (residue L758) and propionyl-CoA (residue Q603), overlaid ^1^H-^15^N HSQC spectra for 250 µM CTD titrated with CoA or propionyl-CoA at a concentration of 0 μM (blue), 62.5 μM (red), 125 μM (green) and 187.5 μM (magenta) with an arrow indicating the direction of chemical shift perturbation during titration. (e) Diffusion coefficient of the CTD in the presence and absence of different CoA analogues. 200 μM proteins with 50 mM sodium phosphate (pD 7.5) in 100% D_2_O were used. Measurements were made at 298 K. Error bars represent standard deviation from different integral regions. (f) Activity of disulfide-stabilised ICL2 mutants compared to wild-type ICL2 in the presence and absence of acetyl-CoA. Error bars represent standard deviation from three measurements.

### Activation of Mtb ICL2 is driven by the binding of CoA activators to the CTD

We then investigated how different CoA analogues modulate the activity of ICL2. Previous work showed that aside from acetyl-CoA, other CoA analogues can also activate ICL2. Among the four CoA analogues tested (acetyl-CoA, propionyl-CoA, succinyl-CoA, CoA), acetyl-CoA was the strongest activator, while CoA was the weakest [10]. We therefore conducted experiments to understand the differences in how acetyl-CoA and CoA interact with the ICL2 CTD and how this is related to the activation of the enzyme.

We first conducted molecular dynamics (MD) simulations of the ICL2 CTD in the presence and absence of acetyl-CoA (based on the ICL2 crystal structures from PDB 6EDW and 6EE1 [10]). In the absence of acetyl-CoA, R665 of the CTD interacts with E640 of its neighbouring CTD, but in the presence of acetyl-CoA, this residue (R665) forms hydrogen bonding interactions with the acetyl group of acetyl-CoA (Table S1-S2). As acetyl-CoA differs from other CoA analogues in the structure of its acetyl moiety, this interaction may strengthen the binding of acetyl-CoA to the CTD.

To investigate whether the difference in the ability of the CoA analogues to activate ICL2 was due to a difference in binding affinity, we measured the binding affinity of the strongest and weakest activators (acetyl-CoA and CoA) to the isolated ICL2 CTD (ICL2_601-766_) using a 1D ^1^H NMR-based assay [15]. The isolated CTD is a suitable model as it shows a high level of structural alignment to the CTDs within full-length ICL2, with r.m.s.d. values of 0.39-0.82 Å (across 163 common Cα positions) as shown by our previous structural work [16]. The binding assay revealed a stronger binding affinity of acetyl-CoA to the isolated CTD, with a *K*_*D*_;value of 39 ± 3 μM, compared to CoA (*K*_*D*_;= 209 ± 9 μM) (Fig. 2b). This was further supported by competitive binding experiments, which showed acetyl-CoA could outcompete CoA for the binding to the CTD (Fig. S2).

The titration experiments above hinted at a relationship between the binding affinity of different CoA analogues and their degree of ICL2 activation observed in our previous study [10], as the weaker binding of CoA and propionyl-CoA to the CTD may have meant that not all ICL2 was bound to the activator and thus not fully activated. Increasing the CoA or propionyl-CoA concentration may therefore help to compensate for its weaker binding by shifting the equilibrium towards the formation of the protein-ligand (CoA-CTD) complex. Indeed, when we tested the activity of ICL2 with a saturating concentration (5 mM) of CoA, we found that it was able to activate the enzyme to an extent similar to acetyl-CoA (Fig. S3). These results suggest that the extent of ICL2 activation correlates to the amount of CTD-activator complex formed in solution, which is likely influenced by the cellular availability of CoA analogues. Given that *Mtb* is unlikely to reach such high intracellular concentrations of CoA analogues, this points to a potential mechanism for regulating ICL2 activity in response to the relative availability or ratio of CoA, acetyl-CoA, and propionyl-CoA.

### Binding of acetyl-CoA promotes the dimerisation of the CTD

Next, we sought to understand the structural changes of the CTD accompanied by acetyl-CoA binding. We used protein NMR spectroscopy, a technique that allows the study of changes to protein dynamics [17], to examine the isolated CTD in the absence and presence of acetyl-CoA. The small size of the isolated CTD (21 kDa) allows us to visualise dynamic events by NMR without the complications that arise from using the full-length protein.

First, we recorded the ^1^H-^15^N heteronuclear single quantum correlation (HSQC) spectra of recombinant ^15^N-labelled CTD (40 µM) in the absence and presence of a saturating concentration of acetyl-CoA (2 mM). The *apo* ^15^N-labelled CTD HSQC spectrum showed a good dispersion of peaks, which indicates that the isolated CTD was folded. Interestingly, upon the addition of acetyl-CoA, most peaks in the resulting HSQC spectrum disappeared (Fig. 2c). This indicates that the CTD may have dimerised (or oligomerised) in the presence of acetyl-CoA, as an increase in molecular weight would reduce the tumbling rate of the protein, which in turn leads to an increased relaxation rate and causes the peaks to broaden [18]. Deuteration of the protein would slow down its relaxation [19] and should therefore lead to better resolution of its peaks in the HSQC spectra. Thus, we produced and purified a partially deuterated form of the CTD (^2^H,^15^N-labelled). With this form, we were able to observe a separate set of peaks that correspond to ^2^H,^15^N-labelled CTD with acetyl-CoA (Fig. S4).

We then conducted titrations of acetyl-CoA and CoA to the ^2^H,^15^N-labelled CTD as this would help determine the exchange rate regime between *apo* and acetyl-CoA/CoA bound-CTD and its dimerisation (or oligomerisation). Titration of acetyl-CoA to the CTD (^2^H, ^15^N-labelled) showed a disappearance of the *apo*-CTD peaks and the appearance of the bound-CTD peaks in the ^1^H-^15^N HSQC spectra (Fig. 2d). This is characteristic of slow exchange (millisecond to second exchange rates). When titrating CoA in place of acetyl-CoA, the exchange between the *apo* and CoA-bound CTD appeared to be fast with a continuous shift of the averaged peak of the two states (picosecond exchange rates). With propionyl-CoA, the *apo* peak gradually shifted to a minimal extent with decreased intensity as the amount of propionyl-CoA increased. This is characteristic of intermediate exchange (nanosecond to microsecond exchange rates). These results confirm our previous findings that acetyl-CoA is the strongest binder to the CTD.

To confirm that the addition of acetyl-CoA resulted in the dimerisation (and not other types of oligomerisation) of the CTD, the isolated CTD was also analysed using diffusion-ordered spectroscopy (DOSY). A standard curve was generated with proteins of known molecular weights (Fig. 2e). The diffusion coefficient of *apo*-CTD was between the coefficients of RNase A (14 kDa) and carbonic anhydrase (29 kDa), suggesting a monomeric protein (21 kDa). The addition of acetyl-CoA increased the molecular weight to around double (Fig. 2e), thus indicating the formation of the CTD dimer. To validate this, a ‘permanent’ CTD dimer was produced by introducing disulfide bonds in the CTD dimer interface by mutating three residues on the interface to cysteines (E640C / N731C / E733C; Fig. S5-S6). The generation of a full-length ICL2 mutant with these mutations is discussed in a later section below (‘fixed bound form’). The molecular weight of the permanent CTD dimer was similar to that of the acetyl-CoA-bound CTD. The DOSY results thus showed that the CTD predominantly exists in the monomeric form in the absence of acetyl-CoA and dimerises in the presence of acetyl-CoA. We also tested the effect of propionyl-CoA and CoA binding on CTD dimerisation by DOSY (Fig. 2e). We found that propionyl-CoA and CoA can also promote full CTD dimerisation at high concentration (1 mM; 5-fold molar excess). Taken together with our kinetic and binding experiments, these results suggest a correlation between acetyl-CoA binding and CTD dimerisation.

### The acetyl-CoA-bound form is the activated form of ICL2

As our results using the isolated CTD indicate a correlation between acetyl-CoA binding and CTD dimerisation, we used full-length ICL2 to investigate how this affects ICL2 activity. Crystal structures of the full-length *apo*- (PDB: 6EDW [10]) and acetyl-CoA-bound ICL2 (PDB: 6EE1 [10]) revealed two distinct CTD dimers with different interfaces as a result of a relative twist between the two monomers. These will be referred to as the *apo*-state and acetyl-CoA-state CTD dimers, respectively. We mutated the residues of full-length ICL2 at the two interfaces to fix the protein in the *apo* and acetyl-CoA bound forms. To create the fixed *apo* forms of full-length ICL2, a triple mutant (ICL2 Q635C / V708C / V734C) was created. For the fixed acetyl-CoA bound form, ICL2 E640C / N731C / E733C was produced (Fig. S5, S7).

Protein mass spectrometry under non-reducing and denaturing conditions was used to determine the masses for both the fixed *apo* and acetyl-CoA-bound forms. The molecular weights of the fixed *apo* and acetyl-CoA bound forms of ICL2 were found to be 175 kDa, which correlates to the expected molecular weight of a dimer (calculated monomeric weight: 87 Da). In contrast, the denatured mass of wild-type ICL2 correlated with the molecular weight of a monomer. Adding the reducing agent tris(2-carboxyethyl)phosphine (TCEP) to the fixed *apo* and fixed acetyl-CoA-bound forms reverted the determined masses to those of monomers (Fig. S8-S10). These results indicate the successful formation of at least one covalent disulfide linkage between dimers, though it is unknown which cysteine pair(s) formed these disulfide bond(s).

To further confirm disulfide-mediated crosslinking of the fixed *apo* form (ICL2 Q635C / V708C / V734C), we crystallised the protein and determined its structure to 2.6 Å resolution (Table S3; PDB: 9OBO). The mutant had a similar overall structure to wild-type ICL2 in the absence of acetyl-CoA (PDB: 6EDW [10]). The fixed *apo* mutant crystallised with a tetramer in the asymmetric unit. Each monomer consists of the N-terminal catalytic domain linked to the C-terminal domain via a flexible linker similar to wild-type ICL2 (PDB: 6EDW [10]). The crystal structure showed electron density for two of the three disulfide bonds for each CTD pair, with S-S distances appropriate for the presence of disulfide bonds for residues Cys(A/B)635 – Cys(C/D)734 and Cys(A/B)734 – Cys(C/D)635 (at ∼ 2.0 Å), but not for Cys(A/B)708 – Cys(C/D)708 (at ∼ 3.7 Å). The modelled positions of the sulfur atoms within these residues were confirmed by calculating anomalous difference Fourier maps using data collected at 8,500 eV (Table S3, Fig. S11). The superposition of the fixed *apo* form crystal structure with that of the wild-type ICL2 structure (PDB: 6EDW [10]) yielded an r.m.s.d. of 4.2 Å for 2920 common Cα positions (Fig. S12). Superposition of the two structures showed the differences between the two structures lie mainly in the positioning of the CTDs (r.m.s.d. of 5.3 Å for 620 common Cα positions of CTDs from four monomeric subunits). The CTDs lie in closer proximity to each other in the fixed *apo* form versus the *apo* ICL2 structure, most likely due to the presence of the introduced disulfide bonds. There was also a slight shift in the unique helical substructure (residues 278-427) (r.m.s.d. of 0.87 Å for 585 common Cα positions from four monomeric subunits). Despite exhaustive attempts, we were unable to crystallise the fixed acetyl-CoA-bound ICL2, but nonetheless, mass spectrometry showed the formation of disulfide bonds in the fixed bound form.

To understand the structure-function relationship of these fixed forms, we characterised their enzyme kinetics (Fig. 2f). We found that the activity of the fixed *apo* form mutant was similar to that of the wild-type ICL2 in the absence of acetyl-CoA, and the addition of acetyl-CoA was unable to activate the fixed *apo* variant. In contrast, the mutant fixed in the acetyl-CoA bound form (ICL2 E640C / N731C / E733C) was found to be as active as wild-type ICL2 in the presence of acetyl-CoA. This indicates that the acetyl-CoA-bound ICL2 form is the activated form of the enzyme regardless of the presence of acetyl-CoA. Hence, the binding of acetyl-CoA (or other CoA analogues) to the CTD converts ICL2 from its unactivated (which has basal activity) to its activated form, but acetyl-CoA itself does not play a role in catalysis.

### NMR and molecular dynamics analysis of acetyl-CoA interactions with the CTD

We then conducted protein NMR experiments to investigate how the binding of acetyl-CoA promotes the formation of the CTD dimer. We first assigned backbone resonances of the isolated CTD in the absence and presence of acetyl-CoA (Table S4–S7). Large chemical shift perturbations were observed between the *apo* and the bound CTD (Fig. 3a). In particular, some of the largest changes in chemical shifts were observed in residues 680-684 and 711-724 (A681, T711, D713, Y716, T718), which form the two α-helices (α25 and α26) that flank the opening of the acetyl-CoA binding pocket (Fig. 3b and Fig. S13). These findings are consistent with the differences observed from the ICL2 crystal structures (PDB: 6EDW and 6EE1 [10]), which show that the binding of acetyl-CoA is accompanied by the movement of these two α-helices away from each other to widen the opening to accommodate the CoA moiety. Many residues with significant chemical shift perturbations belong to the β-hairpin loop between β17 and β18 (F727, T728, V730, Q732, G735, I737, V739, A740), which is located close to α26 and show large structural differences between the *apo* and acetyl-CoA-bound forms (Fig. 3c). Some other residues that showed a large chemical shift difference include those that lie at the dimer interface between the CTDs (Q635, R636, E640, L642, E643, Q663-R665, R668, T669, R674, K701-V705, Y707, V708) (Fig. 3b), as well as residues located close to α25 (S619, L624, K625, L628, and A655). The remaining residues that showed chemical shift perturbations beyond two standard deviations (L601, V606, K609, E613) do not appear to have changes in the crystal structure. However, these residues are solvent-exposed, so they are likely to be sensitive to other factors arising from titration of acetyl-CoA (such as the lithium counter ion).

**Figure 3.**
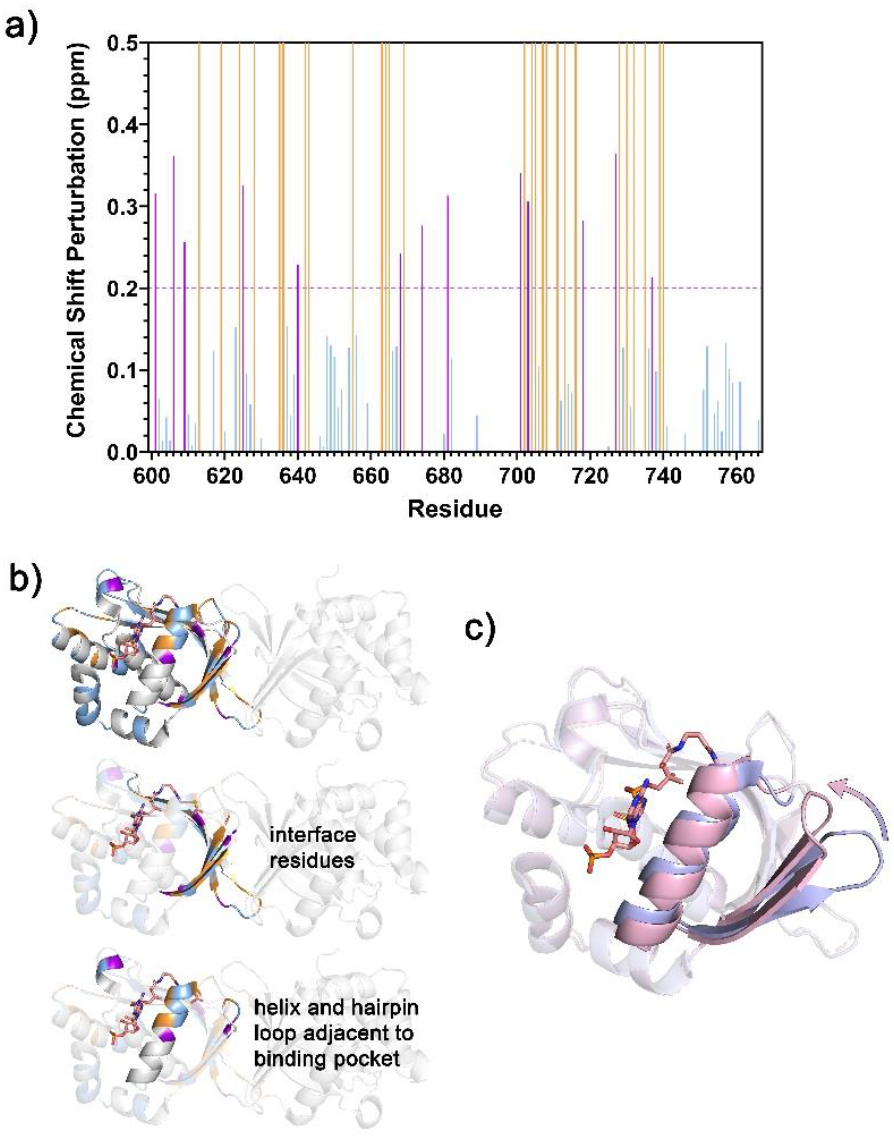
Perturbed residues on the *M. tuberculosis* ICL2 isolated C- terminal domain (CTD) upon acetyl-CoA binding. (a) Chemical shift perturbations (CSPs) between ^1^H and ^15^N resonances in the HSQC spectra of *apo* and acetyl-CoA-bound CTD. The pink line represents two standard deviations (SDs). Residues that were unassigned have zero CSP. Residues that were assigned in either the *apo* or bound spectra are in orange. Residues that had CSPs beyond two SDs are in pink. (b) Residues with significant chemical shift changes mapped onto the CTDs of the acetyl-CoA-bound ICL2 (chains A and C of PDB 6EE1 [10]). Residues are coloured according to (a). The interface residues and the helix and loop adjacent to the acetyl-CoA-binding pocket are highlighted. (c) Overlay of the crystal structures of the CTD in the *apo* (chain A of PDB 6EDW [10]) and acetyl-CoA-bound (chain A of PDB 6EE1 [10]) ICL2. The *apo* form is highlighted in purple, while the bound form is highlighted in pink. The arrow indicates the folding in of the β-hairpin loop. The acetyl-CoA molecule is presented in stick form in pink.

We then conducted ^15^N relaxation and heteronuclear nuclear Overhauser effect (NOE) experiments to study the dynamic nature of these regions. The experiments revealed that the CTD had a longer T1 relaxation time in the presence of acetyl-CoA, which is consistent with the dimerisation of the CTD since a larger size would lead to a slower tumbling rate (Fig. S14). However, aside from the overall increase in T1, there were no changes in the dynamics of the different regions of the CTD. These indicate that the CTD is relatively rigid and that the binding of acetyl-CoA and subsequent dimerization does not affect the rigidity of the domain.

MD simulations were then conducted to examine the conformational changes in the CTD upon acetyl-CoA binding. The simulations were initiated with CTD monomers in the *apo* conformation or the acetyl-CoA-bound conformation with acetyl-CoA either maintained or removed from its binding site. Comparison between MD average conformations of *apo*- and acetyl-CoA-bound CTD revealed changes in backbone positions for residues located on the β-sheets of the CTD, α26 (residues 716-724), and the β-hairpin loop adjacent to this helix (residues 732-734) (Fig. S15). Moreover, when acetyl-CoA was removed from the CTD, residues on α26 were found to move back towards the *apo* conformation (Fig. S16). These results agree with the NMR data, suggesting that acetyl-CoA binding causes α26 to shift outward and the β-hairpin loop to move closer to α26.

### Acetyl-CoA binding selects for the dimer interface in the activated form

MD simulations for the *apo* and acetyl-CoA bound CTD dimers were also conducted, and the results were compared to those for CTD monomers to probe the conformational changes on dimerisation. Comparison of MD average conformations suggested that to establish the CTD dimer in the *apo* state, significant conformational changes were required for residues 663-670 and 731-734, which are located on loops directly involved in the dimer interface (Fig. S17). In contrast, only subtle changes were observed for dimerisation of the CTD with acetyl-CoA bound (Fig. S18). These observations suggest that the acetyl-CoA bound CTD dimers may be more readily formed when the CTDs are in the presence of acetyl-CoA.

Interestingly, one of the loops on the *apo*-state dimer interface (residues 731-734) was found to have significant chemical shift changes in the ^1^H-^15^N HSQC and was identified as flexible in the MD analysis. This loop was shown in the crystal structure to flip away from the interface upon acetyl-CoA binding (Fig. 3c). The flipping of this loop would disfavour the *apo*-state dimer and shift to establish the acetyl-CoA-state dimer. Due to the restriction set by the flexible linker between the N- and C-terminal domains in the full-length ICL2, the twisting motion that accompanies the establishment of the new interface would then cause the CTDs to change their position to the acetyl-CoA-bound form.

### The unique helical substructure is necessary for the oligomeric stability

Finally, we also investigated the role of the unique helical substructure in ICL2 catalysis and activation. To do this, we created an ICL2 mutant (ICL2ΔUHS), with residues 278–428 (belonging to the unique helical substructure) deleted, and residues neighbouring the deletion mutated to match the *Mtb* ICL1 sequence (L273I / L274T / A276E / T277R; Fig. S19). These mutations were made to increase the likelihood of the formation of a folded protein.

CD spectroscopy was first applied to study ICL2ΔUHS. The mutant construct showed the presence of α-helical and β-strand structure elements, implying the formation of a folded protein (Fig. S20). We then subjected ICL2ΔUHS to kinetic analyses. Interestingly, this mutant appeared to have no activity (Fig. 4a). As the unique helical substructure is part of the N-terminal domain that forms the ICL2 tetramer, the deletion of this substructure may affect the oligomeric state of the protein. PISA analysis (“Proteins, Interfaces, Structures and Assemblies” [20]) of the ICL2 crystal structure (PDB: 6EDW) also showed that residues of the unique helical substructure (residues 278-428) form a significant proportion (824 Å^2^) of the total buried surface area (6073 Å^2^) within the dimer interface between chains D/C and chains A/B.

**Figure 4.**
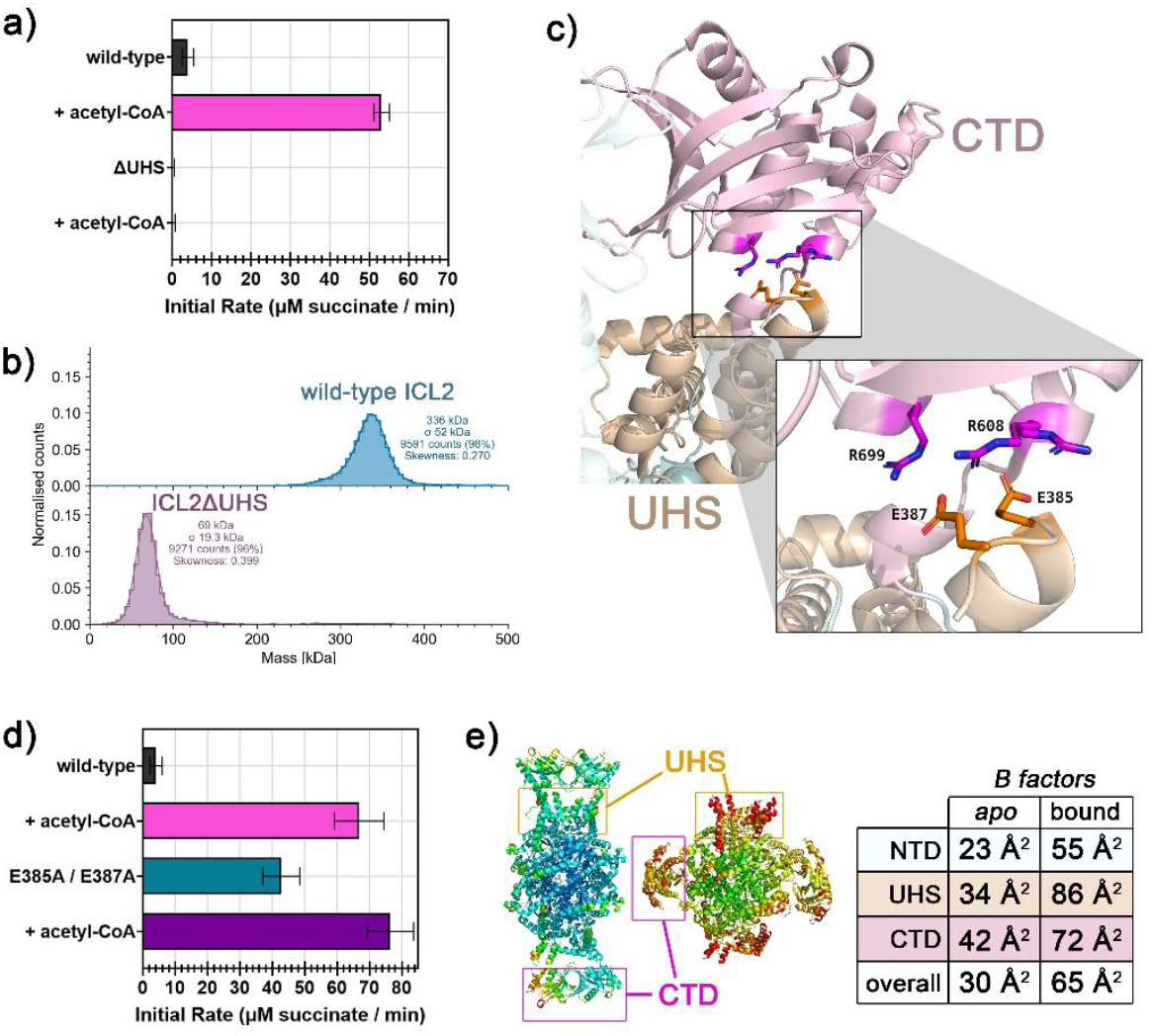
The role of the unique helical substructure (UHS) in *M. tuberculosis* ICL2. Enzyme kinetics measurements were performed at 298 K, and samples contained 500 nM enzyme, 1 mM DL-isocitrate, 5 mM MgCl_2_, 25 μM acetyl-CoA (where applicable), 50 mM Tris-d11 (pH 7.5), 0.02% NaN3 in 90% H_2_O / 10% D_2_O. (a) Effect of UHS deletion on ICL2 activity. Error bars represent standard deviation from three measurements. (b) Mass photometry measurement of ICL2ΔUHS compared to wild-type ICL2. 50 nM protein in 50 mM Tris (pH 7.5) was used. Measurements were made at room temperature. The monomeric molecular weight of wild-type ICL2 and ICL2ΔUHS are 87.5 kDa and 71 kDa, respectively. (c) Interacting residues between the UHS (E385, E387) and the CTD (R608, R699) modelled on the *apo* ICL2 crystal structure (PDB: 6EDW [10]). The UHS is coloured in orange, and the CTD in pink. (d) Activity of the UHS mutant compared to wild-type ICL2 in the presence and absence of acetyl-CoA. Error bars represent standard deviation from three measurements. (e) Average temperature factors of the residues modelled on the crystal structures of *apo* (left; PDB: 6EDW [10]) and acetyl-CoA-bound (right; PDB: 6EE1 [10]) ICL2. The colours of the structures represent the *B* factors of the backbone residues calculated using *B*_average_ from the CCP4 suite. The cartoon is coloured using a rainbow spectrum from blue to red (low to high *B* factor). The UHS and the CTD are highlighted. The average *B* factor in each protein feature is given in a table on the right. NTD stands for the N-terminal domain (excluding UHS).

To investigate the oligomeric state, we performed mass photometry with ICL2ΔUHS. Mass photometry revealed a single peak at around 69 kDa, corresponding to the molecular weight of a monomer. This indicates that the native state of ICL2ΔUHS is a monomer (Fig. 4b). Our results thus showed that the unique helical substructure in ICL2 plays a significant role in the oligomeric stability of ICL2 and that the oligomerisation of ICL2 may be necessary for its activity. This is consistent with previous work with another atypical prokaryotic ICL from *Pseudomonas aeruginosa* ICL that also contains an additional feature within the N-terminal region [21]. Structural superposition of the *P. aeruginosa* ICL (PDB: 6G1O [22]) and *Mtb* ICL2 (PDB: 6EDW [10]) gave an r.m.s.d. of 1.7 Å (over 331 common Cα positions) and showed that this extra segment was inserted in the same region as the unique helical substructure in *Mtb* ICL2 (Fig. S21). However, the number of residues was fewer (25 residues in *P. aeruginosa* ICL vs 151 residues in *Mtb* ICL2). This segment was reported to be important in the oligomeric stability and activity of *P. aeruginosa* ICL [21], in a manner that is similar to our results with *Mtb* ICL2.

### Interactions between the CTD and the unique helical structure mediate activation of ICL2

Careful analysis of the *apo* ICL2 crystal structure (PDB: 6EDW [10]) revealed electrostatic interactions between a flexible loop in the unique helical substructure (E385 and E387) and the CTD (R608 and R699) (Fig. 4c, Table S8-S9). Notably, these interactions were not present in the acetyl-CoA-bound ICL2 (PDB: 6EE1 [10]). These interactions may therefore be important for the kinetics of ICL2. To test this hypothesis, residues from the unique helical substructure were mutated to alanine (ICL2 E385A / E387A). Our results found that ICL2 E385A/E387A was 10x more active than wild-type ICL2 in the absence of acetyl-CoA. Interestingly, upon the addition of acetyl-CoA, the activity of ICL2 E385A/E387A was further increased to the same extent as the wild-type ICL2 in the presence of acetyl-CoA (Fig. 4d). These observations suggest that the interaction between the two protein features via these unique helical substructure residues (E385 / E387) stabilise the CTDs in the unactivated form to prevent the protein from assuming the activated conformation before acetyl-CoA binding. To further support this hypothesis, we conducted a temperature factor analysis of the crystal structures. In the *apo* ICL2 form, the unique helical structure had a similar average *B* factor to the average *B* factor of the full structure, but in the acetyl-CoA-bound ICL2, the unique helical structure had a higher average *B* factor than that of the full structure, indicating that this unique helical substructure is more flexible in the acetyl-CoA-bound ICL2 (Fig. 4e).

## Discussion

*Mtb* ICLs present an interesting and therapeutically relevant system for studying allostery. *Mtb* possesses two isoforms of ICL, one of which can be regulated allosterically by acetyl-CoA [10]. Comparing the structure of the two isoforms revealed two extra features in ICL2, the unique helical substructure and the CTD. Our group previously showed that acetyl-CoA induces drastic conformational changes in ICL2 that involves the repositioning of the CTDs [10]. This study sought to determine the role of each domain in the progression from the binding of acetyl-CoA to the conformational changes that lead to the activation of ICL2.

In the acetyl-CoA-free ICL2 form, the unique helical substructure holds the CTDs in-place (Step 1 in Fig. 5). Structural analyses suggested that the unique helical substructure holds the CTDs in the *apo* state to create a stable interface between CTDs to stabilise the *apo* form before acetyl-CoA binding. Disrupting the interaction between the unique helical substructure and the CTD led to partial activation of the enzyme. Furthermore, the isolated CTD does not appear to form dimers in the absence of acetyl-CoA, so perhaps the unique helical substructure is required to maintain the *apo* state CTD dimers. Acetyl-CoA binding accompanied structural changes in the CTD that involved the folding in of a β-hairpin loop that may disrupt the *apo* CTD dimer interface to favour a new dimer interface (Step 2 in Fig. 5). The conformational changes in this β-hairpin loop were identified from both protein NMR studies and MD simulations. The formation of the new interface, however, is not possible in the *apo* state ICL2 form due to the restrictions set by the linker between the N- and C-terminal domains, leading to the repositioning of the CTDs to the activated form (Step 3 in Fig. 5). Acetyl-CoA then stabilises the new CTD interface in the activated form of the enzyme (Step 4 in Fig. 5). Diffusion coefficient measurements showed that the isolated CTD can form stable dimers in the presence of acetyl-CoA. The dissociation rate of acetyl-CoA from the CTD was also estimated from ZZ-exchange experiments to match with the catalytic rate of ICL2 (koff ∼ 3 s^-1^, Fig. S22), suggesting that acetyl-CoA holds the protein in the activated conformation long enough to establish at least one catalytic cycle.

**Figure 5.**
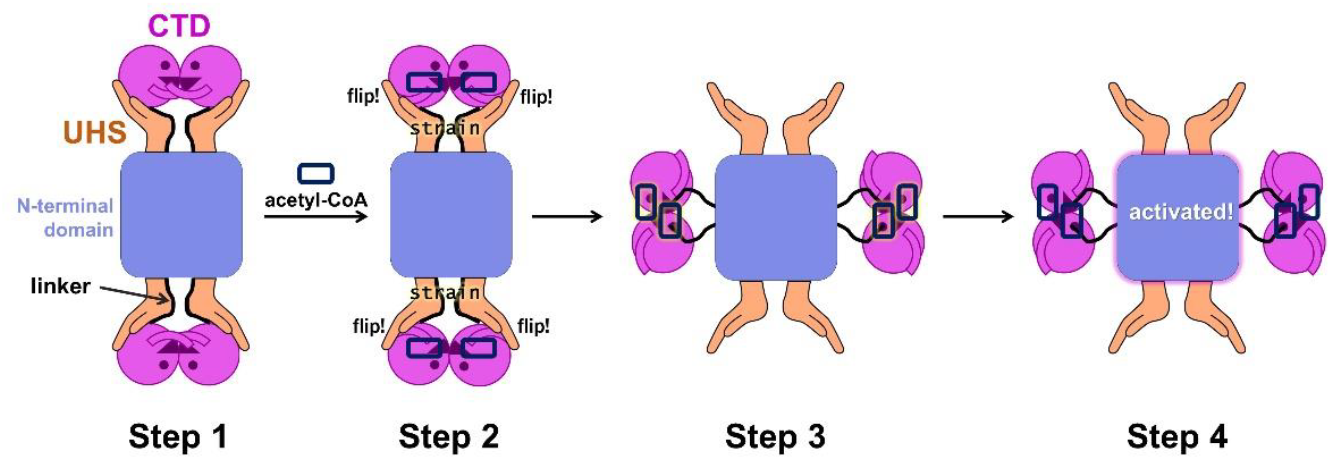
The role of the different structural features in *M. tuberculosis* ICL2 in the journey from acetyl-CoA binding to ICL2 activation. The pink face symbolises the C-terminal domain (CTD) connected to the N-terminal domain (purple square) via a flexible linker (black line). The orange hands represent the unique helical substructure (UHS), and the rounded rectangle with the navy blue outline represents acetyl-CoA. Step 1: The UHS holds the CTD in the unactivated form before acetyl-CoA binding. Step 2: Acetyl-CoA binding leads to the folding in of a β-hairpin loop that is important in the stabilisation of the CTD dimer interface in the unactivated form. Step 3: The destabilisation of the unactivated CTD dimer interface may drive the formation of a new interface that is only possible with the movement of the CTD because of the constraints of the linker. Step 4: The CTD dimers form the new interface that is stabilised by acetyl-CoA, resulting in the activated form of ICL2.

This is the first investigation into the potential role of the unique helical substructure in Group III mycobacterial ICLs. Amino acid networks enable communication between distant sites to link allostery to catalysis [23]. The unique helical substructure appears to be part of this network, but the position of the CTD is still important for the activation, as the removal of the CTD led to the absence of activation. Aside from acetyl-CoA, other CoA analogues can also induce the dimerisation of the CTD and the activation of ICL2. There are many different CoA analogues generated during fatty acid metabolism [24], so it is noteworthy that ICL2 can distinguish between them and preferentially select for acetyl-CoA.

*Mtb* is the causative agent of the lung disease tuberculosis that infected more than 10 million people in 2023. [25]. In most cases, *Mtb* lives inside the unusual niche of alveolar macrophages. Because of this unique environment, *Mtb* has evolved distinct metabolic regulatory mechanisms to aid in its growth and infection [26]. Along with its other isoform, ICL2 was shown to be essential for the growth and virulence of *Mtb* [11]. Enzymes can evolve to alter the metabolic flux within a given organism, leading to novel phenotypes [27]. In the case of *Mtb*, ICL2 has evolved unique structural features that allows its activity to be regulated by metabolites. Although *Mtb* ICLs have long been pursued as targets for tuberculosis drug development, efforts to develop small-molecule inhibitors have been largely unsuccessful due to their compact and highly polar active sites [12]. While the CTD of ICL2 presents an alternative target, its structural similarity to other enzymes such as acetyltransferases poses challenges for developing CTD-specific modulators. The unique structural features and mechanistic insights uncovered in this study, such as the interaction between the unique helical substructure and the CTD, open new possibilities for targeting this enzyme. Our findings may therefore contribute to future drug design strategies aimed at disrupting metabolic flux in *Mtb* as a means to eradicate the pathogen.

## Materials and Methods

### Materials

Unless otherwise stated, all chemicals were purchased from Sigma-Aldrich/Merck, Thermo Fisher Scientific, AK Scientific, and Bio-Rad. Tris-d_11_ and D_2_O were from Cambridge Isotope Laboratories or Cortecnet. Competent cells were obtained from Agilent.

### Recombinant protein production

A synthetic DNA fragment encoding ICL2 of *Mtb* CDC1551 was obtained from Integrated DNA Technologies. The ICL2 gene was subcloned into pYUB28b [28]. pYUB28b and pET28a(+) plasmids encoding genes for ICL2 mutants were synthesised by GenScript. All proteins were synthesised with an N-terminal 6xHis tag.

A detailed methodology for the expression and purification of all recombinant proteins used in this study can be found in the Supplementary Methods. Briefly, recombinant proteins were expressed in either *Escherichia coli* BL21 LOBSTR cells or *E. coli* BL21 LOBSTR cells transformed with the pGro7 plasmid (Takara Bio Inc.) expressing the GroEL/GroES chaperones under the araB promoter. IPTG was added when the cell density reached an OD_600_ of 0.6-0.8 (OD_600_ of ∼10 for deuterated protein expression in ModC1 media). For proteins expressed with the GroEL/GroES chaperones, L-arabinose (final concentration of 0.1% w/v) was added when the cell density reached an OD_600_ of 0.3-0.4. The cells were harvested via centrifugation after overnight incubation at the target induction temperature. The proteins were purified by immobilised metal affinity chromatography (IMAC) and size exclusion chromatography. Purified aliquots were flash-frozen and stored at -80 °C until use.

### Isocitrate lyase activity

NMR data were collected at 298 K using a Bruker 700 MHz AVANCE III HD equipped with a cryoprobe (for testing the kinetics of mutants) or a Bruker 500 MHz AVANCE NEO (for testing CoA vs. acetyl-CoA activation). NMR tubes (5 mm diameter) with a sample volume of 550 µL were used in all experiments. Solutions were buffered using 50 mM Tris-d_11_ (pH 7.5) dissolved in 90% H_2_O and 10% D_2_O. The pulse tip-angle calibration using the single-pulse nutation method (Bruker *pulsecal* routine) was undertaken for each sample [29].

Time course experiments were performed based on previous studies [30]. The reactions were monitored by standard Bruker ^1^H experiments with water suppression by excitation sculpting [31]. The number of transients was 16, and the relaxation delay was 2 s. The lag time between the addition of the enzyme and the end of the first experiment was around 4 min.

### Binding measurement through peak broadening in ^1^H-NMR

The standard Bruker 1D ^1^H *zgesgp* pulse program was used. Peak broadening binding experiments were measured at 298 K on a Bruker 600 MHz AVANCE III HD equipped with a cryoprobe in standard 5 mm NMR tubes. 50 µM acetyl-CoA or CoA was used, and 0-90 (for acetyl-CoA) or 0-250 µM (for CoA) CTD was titrated in. Samples were buffered in 50 mM Tris-d_11_ (pH 7.5) in 90% H_2_O / 10% D_2_O. All spectra were manually processed in TopSpin 4.1.4. Peak broadening was tracked by integrating the singlet CoA proton peak at 8.449 ppm at each titration point and normalised against the *apo* ligand spectrum. Normalised intensities were plotted against protein concentration using non-linear curve fitting by the one-site binding model function on SigmaPlot 15.0.

### ^1^H-^15^N HSQC

^1^H-^15^N HSQC experiments were performed with a Bruker 600 MHz AVANCE III HD, Bruker 700 MHz AVANCE III HD (*apo* ^2^H, ^13^C, ^15^N-CTD), or a Bruker 800 MHz AVANCE III HD (*apo* and acetyl-CoA bound ^2^H, ^13^C, ^15^N-CTD), all equipped with cryoprobes. The size of the FID was 2048 and 256 points for the ^1^H and ^15^N dimension, respectively. The number of scans was 8-16, and the relaxation delay was 1 s. All experiments were conducted at 298 K. For the *apo* and bound ^2^H, ^13^C, ^15^N-CTD, transverse relaxation optimised spectroscopy (TROSY) type HSQC was also collected to increase sensitivity. For the protein assignment, either 750 µM ^13^C, ^15^N-CTD with 2 mM acetyl-CoA (where applicable) or 1.5 mM ^2^H, ^13^C, ^15^N-CTD with 5 mM acetyl-CoA (where applicable) was used. Both were buffered in 50 mM Tris-d_11_ (pH 6.6) in 90% H_2_O / 10% D_2_O.

For the titration of the CTD with the CoA analogues, experiments were conducted at 298 K on a Bruker 500 MHz AVANCE III HD equipped with a cryoprobe. CoA or propionyl-CoA was titrated at concentrations of 0, 62.5, 125, and 187.5 µM to 250 µM ^2^H, ^15^N-CTD. Acetyl-CoA was titrated at concentrations of 0, 50, 100 and 175 µM to 80 µM ^2^H, ^15^N-CTD. Experiments were carried out in 50 mM Tris-d_11_ (pH 6.6) in 90% H_2_O / 10% D_2_O.

### Diffusion ordered spectroscopy (DOSY)

DOSY experiments were carried out at 298 K on a Bruker 500 MHz AVANCE NEO spectrometer in standard 5 mm NMR tubes. 200 µM of the protein sample and 1 mM of the CoA analogue (where applicable) were buffered in 50 mM sodium phosphate (pD 7.5) in 100% D_2_O. The spectra were measured with a quadratic gradient 5-95% in 16 incremental steps, a diffusion delay of 250 ms, a gradient pulse of 1500-2000 µs, and a recycle delay of 10 s. The number of scans was 16. The diffusion coefficients were determined using the T_1_/T_2_ analysis tool from Bruker TopSpin (version 4.1.4).

### Protein backbone assignment

The suite of 3D experiments was undertaken on both the *apo*- and acetyl-CoA-bound CTD: HNCO, HN(CA)CO, HN(CO)CA, HNCA, HN(CO)CACB, HNCACB. Protein backbone assignment was performed using the CCPNMR Analysis software [32]. After the assignments of the *apo* and bound datasets, the assignments from the conventional HSQC spectrum of the *apo* CTD was transferred to the TROSY HSQC spectrum, then the minimal shift assumption resonance translation method [33] was used to transfer the peaks from the *apo* CTD TROSY HSQC spectrum to the unassigned peaks in the bound CTD TROSY HSQC spectrum or vice versa. After the transfer of assignment, the CO, Cα, and Cβ peaks in the transferred peak was checked to see if the assignment of the peak is consistent. Peaks that could not be unambiguously transferred were removed from the analysis.

### Molecular dynamics simulations

MD simulations were conducted using NAMD 2.13 [34], and trajectories were visualised and analysed in VMD [35]. MD simulations were set up for CTD monomers and dimers starting from conformations extracted from crystal structures of *apo* ICL2 (PDB 6EDW [10]) and acetyl-CoA bound ICL2 (PDB 6EE1 [10]). For the CTD monomer, additional MD simulations were set up starting from the acetyl-CoA bound conformation but with acetyl-CoA removed. Initial force field topology and parameters for acetyl-CoA were obtained from CGenFF server (https://cgenff.paramchem.org), from which parameters were assigned by analogy to existing parameters in the force field [36-39]. A penalty score is given to each assigned parameter and those with large penalty values were further refined by Force Field Toolkit [40]. Explicit TIP3 water molecules were added to solvate the protein molecules in a water box in VMD. Na^+^ and Cl^-^ ions were added to balance the net charge of the water box. MD simulations were conducted with CHARMM36 force field [41] at a constant temperature and pressure (310 K, 1 atm). The cutoff distance for van der Waals interactions was set to 12 Å. In each simulation, the system was first minimised for 5000 steps followed by dynamics simulation conducted with 2 fs time steps. Three MD simulations were conducted for each MD system, initiated with different random seeds (Table S10). Trajectory frames were collected every 100 ps. Trajectories from equilibrated time period of the MD simulations as indicated by the protein backbone r.m.s.d. values (Fig. S23) were used in analysis.

## Acknowledgments

We thank Dr Biswaranjan Mohanty for his assistance and advice with the 3D NMR experiments for the labelled C-terminal domain that was carried out at Sydney Analytical, a core research facility at the University of Sydney. We also thank Dr Michael Schmitz from the Nuclear Magnetic Resonance Centre at the University of Auckland for his expertise. We acknowledge Dr Richard Hopkinson (University of Leicester) for his helpful comments and discussions on earlier drafts of this paper. We acknowledge the support of the Australian Government in provision of access to ANSTO’s National Deuteration Facility, which is partly funded through the National Collaborative Research Infrastructure Strategy (NCRIS), via NDF proposal 15636 for the generation of the deuterated ICL2 C-terminal domain. We would also like to acknowledge the use of the Magnetic Resonance Facility, Melbourne Protein Characterisation, and the Mass Spectrometry and Proteomics Facility at the Bio21 Molecular Biology and Biotechnology Institute for this work. NAMD was developed by the Theoretical and Computational Biophysics Group in the Beckman Institute for Advanced Science and Technology at the University of Illinois at Urbana-Champaign. The authors wish to acknowledge the contribution of NeSI (https://www.nesi.org.nz) high performance computing facilities to the results of this research. NZ’s national facilities are provided by the NZ eScience Infrastructure and funded jointly by NeSI’s collaborator institutions and through the Ministry of Business, Innovation & Employment’s Research Infrastructure programme. Part of this study was conducted using the MX2 beamline at the Australian Synchrotron, which is part of ANSTO and made use of the ACRF detector. We thank the beamline staff for their enthusiastic and professional support.

E.Y.W.H. is supported by the Melbourne Research Scholarship, Rowden White Scholarship, Norma Hilda Scholarship, Dame Margaret Blackwood Soroptimist Scholarship, and Dr Albert Shimmins Postgraduate Writing-Up Award from the University of Melbourne. B.X.C.K. was supported by the University of Auckland Doctoral Scholarship. G.B., J.T., W.J., and I.K.H.L. are supported by the Marsden Fund (21-UOA-108) by the Royal Society of New Zealand. G.B. and I.K.H.L. are also supported by the Project Grant from the Maurice & Phyllis Paykel Trust. J.T. is also supported by the a Ngā Puanga Pūtaiao Fellowship through the Royal Society of New Zealand. M.J.M was funded by an Australian Research Council (ARC) Future Fellowship (FT180100397). I.K.H.L. and M.J.M. would like to thank the University of Melbourne for support through the Driving Research Momentum (DRM) initiative.

