## Supplementary Information for "Dynamic allostery drives acetyl-CoA-mediated activation of *Mycobacterium tuberculosis* isocitrate lyase 2"

This PDF file includes:

Supplementary Methods
Figures S1 to S22
Tables S1 to S10
SI References

**Supplementary Methods**

***Recombinant protein expression***

| Protein | Plasmid Backbone | Expression System (*E.coli* strain) | Media | [IPTG] | Induction Temp. |
|---|---|---|---|---|---|
| ICL2 | pYUB28b | BL21 LOBSTR (+pGro7) | terrific broth (TB) | 0 mM (leaky) | 37 °C |
| ICL2 fixed *apo* form (Q635C/V708C/V734C) | pYUB28b | BL21 LOBSTR (+pGro7) | terrific broth (TB) | 0 mM (leaky) | 37 °C |
| ICL2 fixed bound form (Q635C/V708C/V734C) | pYUB28b | BL21 LOBSTR (+pGro7) | terrific broth (TB) | 0 mM (leaky) | 37 °C |
| ICL2 CTD ($ICL2_{601-766}$) | pYUB28b | BL21 LOBSTR | TB auto-induction | N/A | 37 °C |
| $^{15}N$-CTD | pYUB28b | BL21 LOBSTR | M9 minimal media | 1 mM | 28 °C |
| $^{2}H$, $^{15}N$-CTD (HSQC, Relaxation) | pYUB28b | BL21 LOBSTR | M9 minimal media | 1 mM | 28 °C |
| $^{13}C$, $^{15}N$-CTD | pYUB28b | BL21 LOBSTR | M9 minimal media | 1 mM | 28 °C |
| $^{2}H$, $^{15}N$-CTD (ZZ-exchange) | pET28a(+) | BL21 LOBSTR | ModC1 minimal media | 1mM | 20 °C |
| $^{2}H$, $^{15}N$, $^{13}C$-CTD | pET28a(+) | BL21 LOBSTR | ModC1 minimal media | 1mM | 20 °C |
| ICL2 E385A/E387A | pYUB28b | BL21 LOBSTR (+pGro7) | terrific broth (TB) | 0 mM (leaky) | 37 °C |
| ICL2ΔUHS | pYUB28b | BL21 LOBSTR (+pGro7) | terrific broth (TB) | 0.5 mM | 18 °C |

For proteins expressed in the pYUB28b plasmid backbone, 50 µg/mL hygromycin was added, and for those co-transformed with pGro7, 34 µg/mL chloramphenicol was also added. To produce $^{15}N$-labelled, $^{2}H$, $^{15}N$-labelled (for HSQC and relaxation experiments), and $^{13}C$, $^{15}N$-labelled CTD, the components in M9 minimal media were replaced with their respective labelled variants to achieve the desired labelling. For $^{15}N$ labelling, $^{15}NH_4Cl$ was used. For $^{2}H$, $^{15}N$ labelling, $^{15}NH_4Cl$ and $D_2O$ (99.8 atom %D) were used. The average deuteration for this protein was 63 %D. For $^{13}C$, $^{15}N$ labelling, $^{15}NH_4Cl$, $D_2O$ (99.8 atom %D), and $^{13}C$-glucose (5 g/L) were used.

For proteins expressed in the pET28a(+) backbone, 40 µg/mL kanamycin was added to the ModC1 minimal media. To produce $^{2}H$,$^{15}N$-labelled (for ZZ-exchange) and $^{2}H$, $^{13}C$, $^{15}N$ -labelled CTD, a previously described high-cell density protocol in ModC1 minimal media was used [1]. The components in ModC1 minimal media were replaced with their respective labelled variants to achieve the desired labelling. For $^{2}H$, $^{15}N$ labelling, $^{15}NH_4Cl$, $D_2O$ (99.8 atom %D), and glycerol-$d_8$ (20 g/L) were used. For $^{2}H$, $^{13}C$, $^{15}N$ labelling $^{15}NH_4Cl$, $D_2O$ (99.8 atom %D), and $^{13}C$-glycerol (20 g/L) were used. The average deuteration (non-exchangeable hydrogen atom positions) of the expressed double and triple-labelled CTD were 98 and 84 %D respectively.

### *Protein purification*

| | Buffer A | Buffer B | Storage Buffer |
|---|---|---|---|
| ICL2 and mutants (except ICL2$_{601\text{-}766}$) | 50 mM HEPES<br>150 mM NaCl<br>5 mM imidazole<br>1 mM β-mercaptoethanol<br>pH 7.5 | 50 mM HEPES<br>150 mM NaCl<br>500 mM imidazole<br>1 mM β-mercaptoethanol<br>pH 7.5 | 50 mM HEPES<br>150 mM NaCl<br>pH 7.5 |
| CTD / ICL2$_{601\text{-}766}$ (including labelled versions) | 50 mM HEPES<br>150 mM NaCl<br>5 mM imidazole<br>pH 7.5 | 50 mM HEPES<br>150 mM NaCl<br>500 mM imidazole<br>pH 7.5 | 50 mM HEPES<br>150 mM NaCl<br>pH 7.5 |

The cells containing the protein of interest were lysed by sonication, then the cell lysate was clarified via centrifugation. For immobilised metal affinity chromatography (IMAC), either the HisTrap HP (5 mL) or HisGraviTrap (Cytiva) column was used. The column was charged with 5 column volumes of 50 mM $NiSO_4$ and equilibrated with Buffer A before applying the filtered cell lysate. 15-30 column volumes of a 1:9 ratio of Buffer A:B was used to wash out non-specifically bound proteins prior to elution with Buffer B. Fractions containing the target protein were combined and dialysed overnight at 4 °C using a dialysis tubing (Thermo Fisher) into the Storage Buffer. After IMAC, the proteins (except for the labelled proteins) were purified further via size exclusion chromatography using the HiLoad 16/600 Superdex 75 pg (Cytiva) column. Storage Buffer was used for elution.

### *Protein backbone assignment*

For the $^{13}C$, $^{15}N$ *apo* CTD, the 3D spectra were recorded at 298 K on a Bruker 600 MHz AVANCE III HD equipped with a cryoprobe. Standard 5 mm NMR tubes with a sample volume of 500 µL were used. The sample contained 750 µM $^{13}C$, $^{15}N$-CTD buffered in 50 mM Tris-$d_{11}$ (pH 6.6) in 90% $H_2O$ / 10% $D_2O$. The acquired data were zero filled on Bruker TopSpin (version 4.1.4).

The 3D spectra for the acetyl-CoA-bound CTD were recorded at 298 K on a Bruker 800 MHz AVANCE III HD equipped with a cryoprobe. A 3 mm Shigemi tube with a sample volume of 120 µL was used. The sample contained 1.5 mM $^{2}H$, $^{13}C$, $^{15}N$-labelled CTD, 5 mM acetyl-CoA, 50 mM Tris-$d_{11}$ (pH 6.6), 0.02% $NaN_3$ in 90% $H_2O$ / 10% $D_2O$. These spectra were recorded using 10% non-uniform sampling (NUS) and Poisson gap sampler [2]. Spectra were reconstructed with the compressed sensing algorithm using qMDD [3] and processed using NMRPipe [4].

$^{15}N$ longitudinal relaxation ($T_1$), $^{15}N$ transverse relaxation ($T_2$), and $^{15}N$-$^{1}H$ nuclear Overhauser enhancement (NOE) experiments were undertaken using a Bruker 600 MHz AVANCE III HD spectrometer equipped with a cryoprobe at 298 K in 5 mm standard NMR tubes. 300 µM of $^{2}H$, $^{15}N$-CTD with or without 2 mM acetyl-CoA buffered in 50 mM Tris-$d_{11}$ (pH 6.6) in 90% $H_2O$ / 10% $D_2O$ was used. For $T_1$ experiments, the measurements were performed with 8 scans, a relaxation delay of 6 s, and 2048 and 256 points for the $^{1}H$ and $^{15}N$ dimension, respectively. Nine delay points of 0.02, 0.10, 0.20, 0.40, 0.70, 1.10, 1.60, 2.20, and 2.90 s with two duplicates were recorded. For $T_2$ experiments, the measurements were performed with 8 scans, a relaxation delay of 6 s, and 2048 and 512 points for the $^{1}H$ and $^{15}N$ dimension, respectively. Nine delay points of 1, 2, 3, 5, 7, 9, 11, 14, and 17 ms with two duplicates were recorded. For NOE experiments, two spectra with or without $^{1}H$ saturation were recorded with 16 scans, a relaxation delay of 5 s, and 2048 and 512 points for the $^{1}H$ and $^{15}N$ dimension, respectively. All acquired data were zero-filled on Bruker Topspin (version 4.1.4) before analysed using the Relaxation Analysis module in the CCPNMR Analysis software [5]. The fitting of the curves (for $T_1$/$T_2$) and the peaks were checked, and residues with poor fitting or poor signal-to-noise ratio were removed from the analysis.

***Circular dichroism (CD) spectroscopy***

CD spectra were collected with Applied Photophysics Chirascan plus spectrometer at 25 °C with a 0.2-cm pathlength cell. The protein sample was diluted to 0.1 mg/mL with 10 mM sodium phosphate (pH 7.5), then the spectra were obtained from 190 nm to 260 nm with scanning speed of 120 nm/min and 1 nm bandwidth. The CD spectrum of the protein was subtracted from the CD spectrum of the buffer-only blank.

***Mass spectrometry***

LC-MS was carried out using the Agilent 6520 Accurate-Mass Q-TOF. The LC system was equipped with the Jupiter® 5 µm column (Phenomenex, C5, 300 Å, 50 x 2 mm). Proteins (1-5 µg) were injected into the column at an isocratic flow of 250 µL/min. The eluents were 0.1% (v/v) formic acid in water (solvent A) and 95% (v/v) acetonitrile containing 0.1% (v/v) formic acid in water (solvent B). The flow gradient was (i) 0-5 min at 5% B; (ii) 5-13 min, 5-95% B; (iii) 13-14 min, 95% B; (iv) 14-15 min, 95-5% B. A wash was done between samples by running the program with 10 µL trifluoroethanol (Acros Organics). The mass spectra were acquired in positive mode with dual electrospray ionization (ESI) voltage of 4 kV with MS spectra scanning from m/z 100-3200 at a rate of 2 spectra/s. Data analysis was done using the MassHunter Quanlitative Analysis Software (Agilent). The isotope series in the obtained mass spectra was deconvoluted at a mass range of 10-200 kDa with a 1 Da mass step and proton as an adduct to determine the intact molecular weight of the denatured protein.

***Mass photometry***

Mass photometry measurements were performed using a Refeyn Two$^{MP}$ mass photometer. A standard curve was created using bovine serum albumin (66.5 kDa monomer and 132 kDa dimer) and thyroglobulin (670 kDa). 50 nM protein samples in 50 mM Tris (pH 7.5) were used. Data was collected at room temperature for 60 s and analysed using the DiscoverMP software (Refeyn).

***Crystallisation of the ICL2 disulfied-stabilised mutants***

The purified mutants were supplemented with a final concentration of 1 mM $MgCl_2$ and 1 mM succinate and diluted to the protein concentration required for the screening with 50 mM HEPES (pH 7.5), 150 mM NaCl. Initial crystallisation trials were conducted using the commercially available screens Index (Hampton Research) and Morpheus (Molecular Dimensions) by sitting drop vapour diffusion in 96-well plates (Molecular Dimensions) at two different protein concentrations (4 and 8 mg/mL). Crystallisation drops consisting of equal volumes (125 nL) of reservoir and protein solutions were dispensed using a mosquito® Xtal3 (SPT Labtech) and were equilibrated against a 50-µL reservoir of screen solution. Plates were incubated at 20 °C. Fixed *apo* form crystals were obtained after 7 days in conditions B10 (0.1 M Tris/Bicine, pH 8.5, 20% v/v ethylene glycol, 10% w/v PEG8000, 30 mM each of sodium fluoride, sodium bromide, and sodium iodide) and H10 (0.1 M Tris/Bicine, pH 8.5, 20% v/v ethylene glycol, 10% w/v PEG8000, 20 mM each of DL-glutamic acid monohydrate, DL-alanine, glycine, DL-lysine monohydrochloride, and DL-serine) of the Morpheus screen. No crystals were observed for the fixed bound form.

Optimisation of the Morpheus condition B10 was carried out by hanging drop vapour diffusion in 24-well VDX plates (Hampton Research) with drops containing equal volumes (1 µL) of protein (4 mg/mL protein with 1 mM $MgCl_2$ and 1 mM succinate in 50 mM HEPES, pH 7.5, 150 mM NaCl) and crystallisation solution equilibrated against 500 µL reservoir solution. Diffraction-quality crystals of the fixed *apo* mutant grew after 9 days at 20 °C in one of the wells (0.1 M Tris/Bicine, pH 8.5,

20% ethylene glycol, 10% w/v PEG8000) to 300 x 60 x 40 µM. For data collection, the crystals were flash-cooled in liquid nitrogen directly from the crystallization drop.

Diffraction data that were used to solve and refine the structure were recorded on an Eiger 16M detector at the Australian Synchrotron on beamline MX2 at a wavelength of 0.954 Å. To confirm the positions of the sulfur atoms within the model, anomalous difference Fourier maps were calculated from datasets collected at 8,500 eV. All data were collected at 100 K and were processed with XDS [6] and merged and scaled with AIMLESS [7] (Table S5).

The crystal structure of the fixed *apo* mutant was determined by molecular replacement with PHASER [8] from the CCP4 suite [9], using the coordinates of the full-length *apo* ICL2 (PDB: 6EDW) (Bhusal et al, 2019) as a search model (after the removal of all water molecules). Model building and the addition of water molecules were carried out in COOT [10] and model refinement with REFMAC5 [11] with TLS [12]. The quality of the structure was determined by MolProbity [13]. Data collection and refinement statistics are detailed in Table S5.

Structure superpositions and the calculation of r.m.s.d's were carried out with LSQKAB from the CCP4 suite [9], and analyses of interactions between monomers were performed with the PDBePISA server [14].

### *ZZ-exchange*

ZZ-exchange experiments for the analysis of chemical exchange between the monomer and dimer states of the CTD were conducted at 298 K using Bruker 500 MHz AVANCE III HD equipped with a cryoprobe in standard 5 mm NMR tubes. 300 µM of $^2$H, $^{15}$N-CTD buffered in 50 mM Tris-$d_{11}$ (pH 6.6) in 90% $H_2O$ / 10% $D_2O$ was used. 500 µM of acetyl-CoA was titrated into the protein to reach a 1:1 monomer:dimer ratio as judged by the ratio of monomer:dimer S650 peak in the $^1$H-$^{15}$N HSQC. Two dimensional ZZ-exchange spectra were measured with 2048 and 512 points in the $^1$H and $^{15}$N dimensions, respectively. The number of scans was 8. A series of spectra were acquired at mixing times of 25, 50, 75, 100, 150, 200, 250, and 400 ms. The peak intensities of the monomer and dimer states of residues S650 ($^1$H and $^{15}$N peak of monomer: 8.68 ppm, 114.5 ppm; dimer: 8.75 ppm, 114.7 ppm) and Q623 ($^1$H and $^{15}$N peak of monomer: 7.48 ppm, 115.3 ppm; dimer: 7.40 ppm, 115.0 ppm) were obtained using Bruker Topspin (version 4.1.4) and plotted using Excel. If a peak could not be unambiguously identified for any of the states at a certain timepoint, that timepoint was omitted from the analysis. The equation below was used to calculate the $k_{on}$ and $k_{off}$ rate.

$$\boldsymbol{\Xi(t)} = \frac{\boldsymbol{a_{DM}(t)a_{MD}(t)}}{\boldsymbol{a_{MM}(t)a_{DD}(t) - a_{DM}(t)a_{MD}(t)}} \cong \boldsymbol{\zeta t^2 = 4k_{off}k_{on}[M]t^2} = \frac{\boldsymbol{4k_{off}}^{\mathbf{2}}}{\boldsymbol{K_D}}\boldsymbol{[M]t^2}$$

**Equation 1. The relationship between the composite ratio of auto and cross peaks and the exchange rate in ZZ-exchange.** $a_{XY}(t)$ represents the resonance intensity arising from state X and detected on state Y. D represents the dimer, and M represents the monomer. $\boldsymbol{\zeta}$ is obtained by fitting a quadratic equation using Excel 2016. $k_{off}$ and $k_{on}$ represents the kinetic off and on rate, respectively. $K_D$ represents the dissociation constant obtained from the binding experiment, and [M] is the equilibrium monomer concentration.

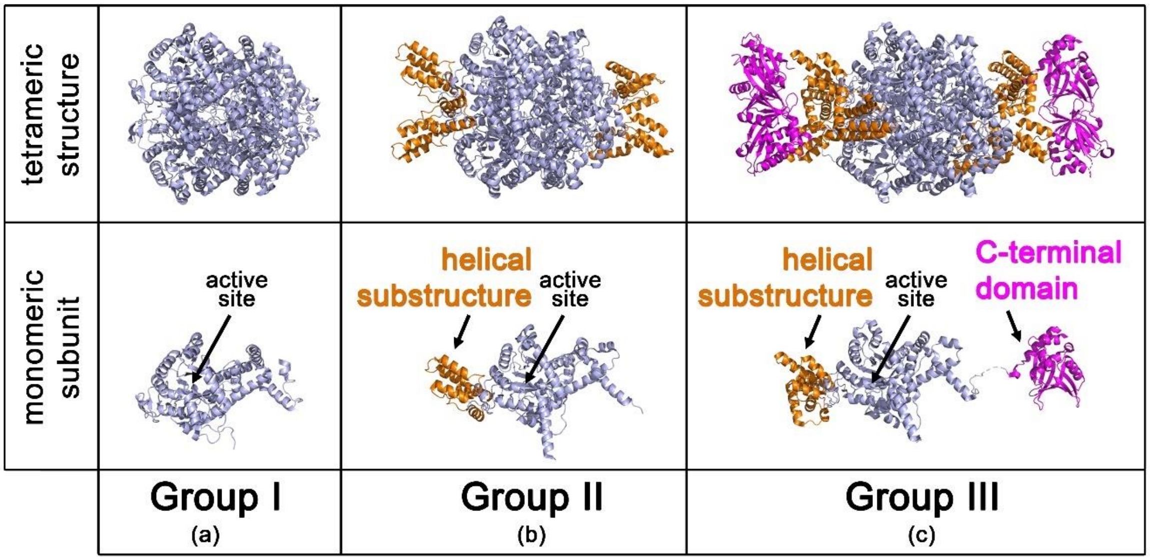


**Figure S1. Comparison of different groups of isocitrate lyases.** The top row illustrates the tetrameric form, while the bottom row shows one monomeric chain from each of the tetramers. The unique domains in each group are highlighted. (a) Group I ICL illustrated by *Mycobacterium tuberculosis* isocitrate lyase 1 (PDB: 5DQL [15]). (b) Group II ICL illustrated by *Saccharomyces cerevisiae* isocitrate lyase (PDB: 7EBC [16]). The unique helical substructure is coloured in orange. (c) Group III ICL illustrated by *Mycobacterium tuberculosis* isocitrate lyase 2 (PDB: 6EDW [17]). The unique helical substructure is coloured in orange and the C-terminal domain in magenta.

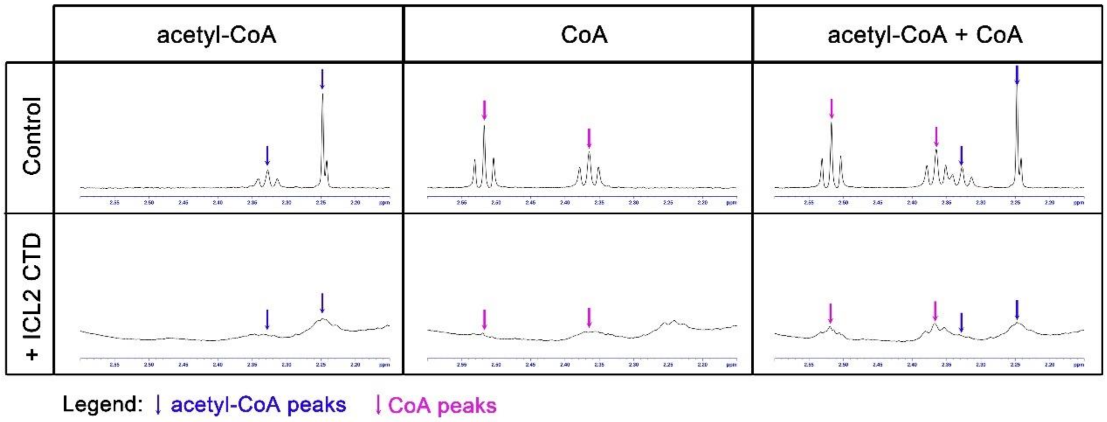


**Figure S2. Binding of a mixture of acetyl-CoA and CoA to the isolated C-terminal domain (CTD).** Measurements were performed at 298 K, and samples contained 50 µM CTD (where applicable), 50 µM acetyl-CoA and/or 50 µM CoA, 50 mM Tris-$d_{11}$ (pH 7.5), 0.02% $NaN_3$ in 90% $H_2O$ / 10% $D_2O$. The top row contains the spectra of the CoA analogue alone, while the bottom row shows the spectra in the presence of the CTD. The blue arrows indicate the acetyl-CoA peaks, and the pink arrows indicate the CoA peaks.

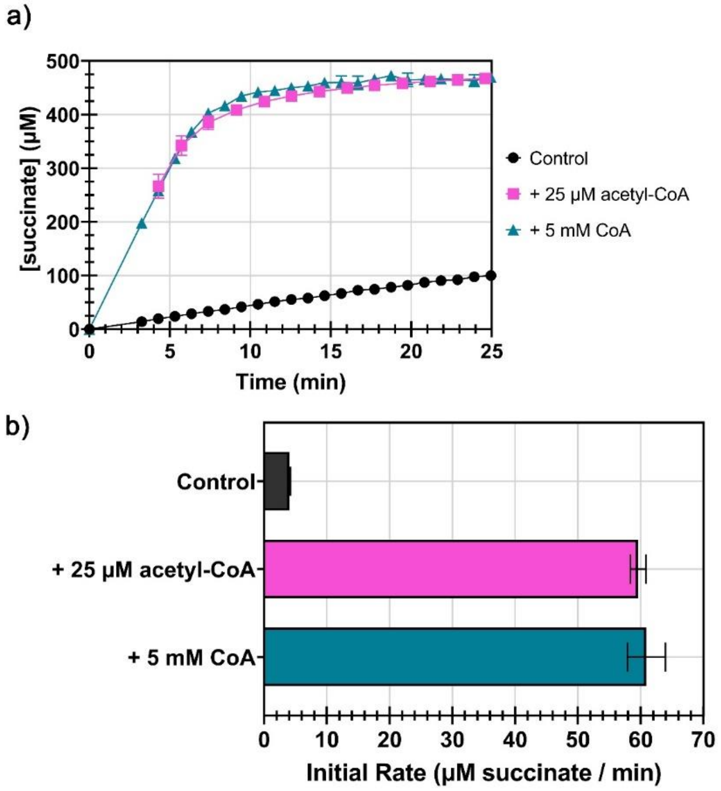


**Figure S3. Activity of ICL2 in the presence of acetyl-CoA or excess CoA.** Measurements were performed at 298 K, and samples contained 500 nM enzyme, 1 mM DL-isocitrate, 5 mM $MgCl_2$, 25 μM acetyl-CoA or 5 mM CoA (where applicable), 50 mM Tris-$d_{11}$ (pH 7.5), 0.02% $NaN_3$ in 90% $H_2O$ / 10% $D_2O$. Error bars represent standard deviation from three measurements. (a) Time course of the reaction as illustrated by the formation of the product succinate over time. (b) The rate of the reaction expressed as the amount of succinate formation per minute.

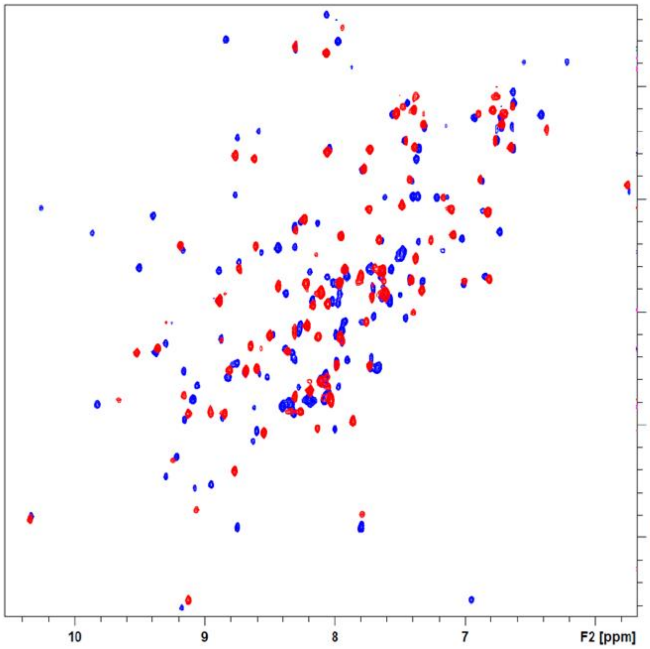


**Figure S4. Effect of acetyl-CoA binding on $^{2}H,^{15}N$-labelled ICL2 C-terminal domain (CTD).** Overlay of $^{1}H$-$^{15}N$ HSQC spectra of 40 μM of $^{2}H,^{15}N$-CTD in the absence (blue) and presence (red) of 2 mM acetyl-CoA. The number of peaks in the HSQC spectrum of the acetyl-CoA-bound ICL2 C-terminal domain improved significantly with deuteration (Refer to Fig. 2c for HSQC spectrum of non-deuterated protein).

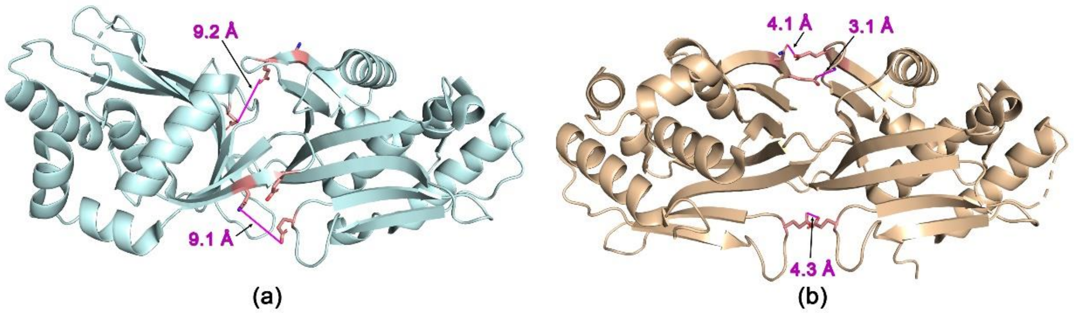


**Figure S5. Modelling the mutated residues of the fixed acetyl-CoA-bound form (E640C/N731C/E733C) on the ICL2 C-terminal domains (CTDs).** The numbers shown are the distances (in Å) of the closest among the three mutated residues between the two CTDs. (a) The three mutated residues on the CTDs of *apo* ICL2 (PDB: 6EDW [17], chain D on the left and A on the right). From top to bottom: E640-E733, E733-E640. N731 was facing away from the interface, making an interaction with the opposing CTD unlikely, hence the distance was not measured. (b) The three mutated residues on the CTDs of acetyl-CoA-bound ICL2 (PDB: 6EE1 [17], chain C on the left and A on the right). From top to bottom: N731-E733, E733-N731, E640-E640.

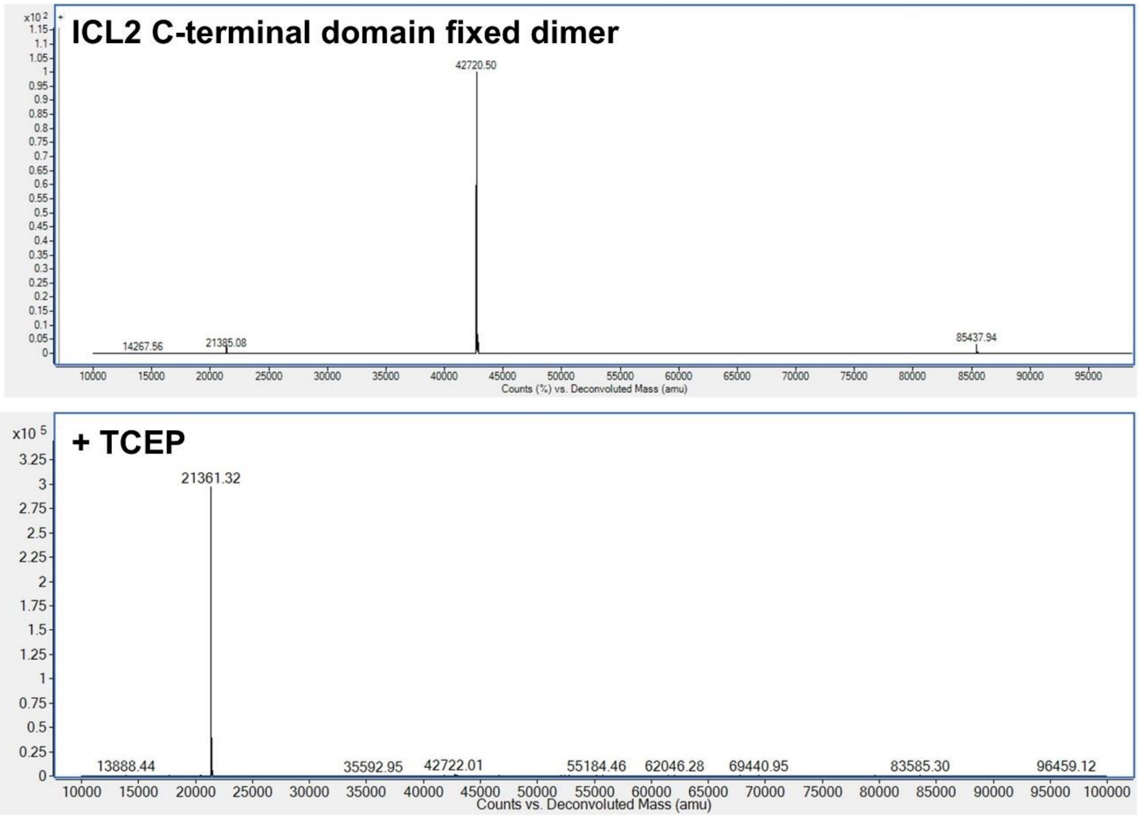


**Figure S6. Deconvolution spectra of the isolated C-terminal domain (CTD) of ICL2 fixed acetyl-CoA-bound form (ICL2$_{601\text{-}766}$ E640C/N731C/E733C) obtained via denaturing mass spectrometry.** The top spectrum represents the protein on its own, while the bottom spectrum represents the protein in the presence of TCEP. The average mass of the ICL2 fixed bound form CTD is 21493 Da, while the observed mass was 21361 Da. The mass loss of around 131 Da may correspond to the loss of the N-terminal methionine. Under denaturing conditions, the protein appears to form a covalent dimer (~42 kDa) that can be reduced to a monomer (~21 kDa) in the presence of TCEP. The calculated mass of a dimer (no N-terminal methionine) with three disulfide bonds (-2 Da for each disulfide bond formation) was 42718 Da, and the observed mass was 42721 Da.

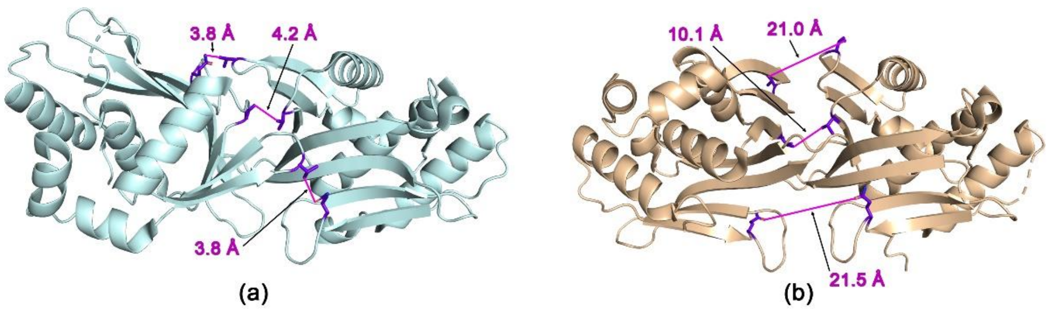


**Figure S7. Modelling the mutated residues of the fixed *apo* form (Q635C/V708C/V734C) on the ICL2 C-terminal domains (CTDs).** The numbers shown are the distances (in Å) of the closest among the three mutated residues between the two CTDs. (a) The three mutated residues on the CTDs of *apo* ICL2 (PDB: 6EDW [17], chain D on the left and A on the right). From top to bottom: Q635-V734, V708-V708, V734-Q635. (b) The three mutated residues on the CTDs of acetyl-CoA-bound ICL2 (PDB: 6EE1 [17], chain C on the left and A on the right). From top to bottom: V734-V734, V708-V708, Q635-Q635.

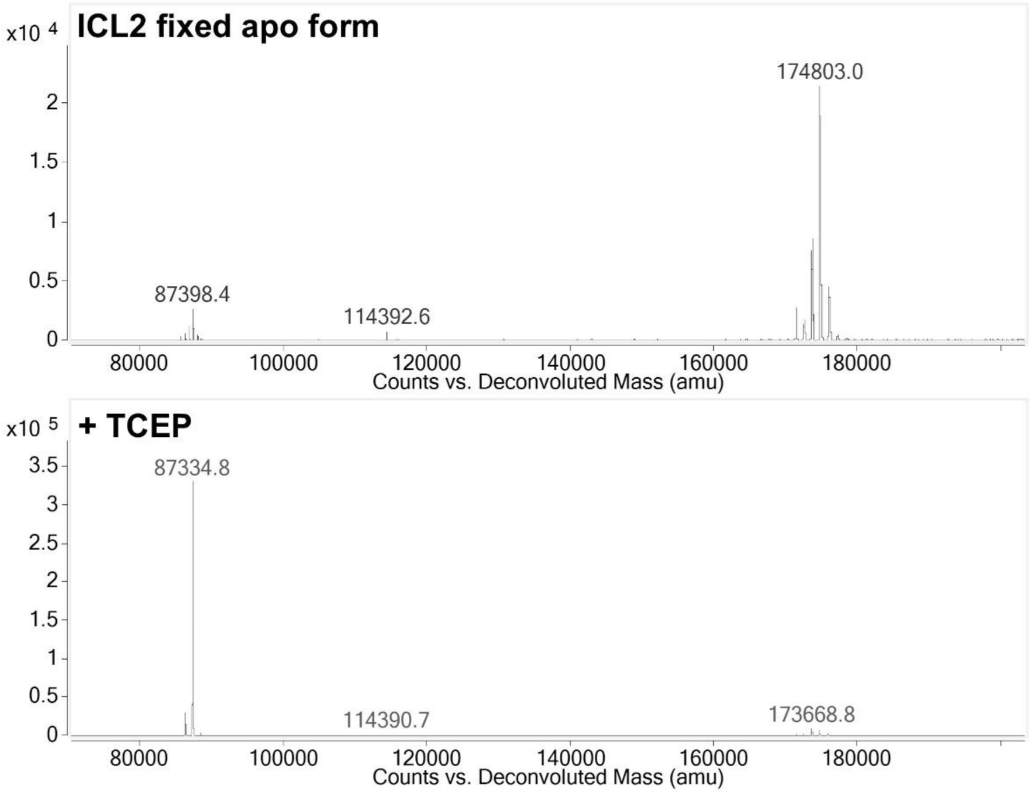


**Figure S8. Deconvolution spectra of the ICL2 fixed *apo* form (Q635C/V708C/V734C) obtained via denaturing mass spectrometry.** The top spectrum represents the protein on its own, while the bottom spectrum represents the protein in the presence of TCEP. The average mass of the ICL2 fixed *apo* form is 87466 Da, while the observed mass was 87335 Da. The mass loss of 131 Da may correspond to the loss of the N-terminal methionine. Under denaturing conditions, the protein appears to form a covalent dimer (~170 kDa) that can be reduced to a monomer (~87 kDa) in the presence of TCEP. The calculated mass of the ICL2 fixed *apo* form dimer (no N-terminal methionine) with three disulfide bonds (-2 Da for each disulfide bond formation) is 174664 Da, and the observed mass is 174803 Da.

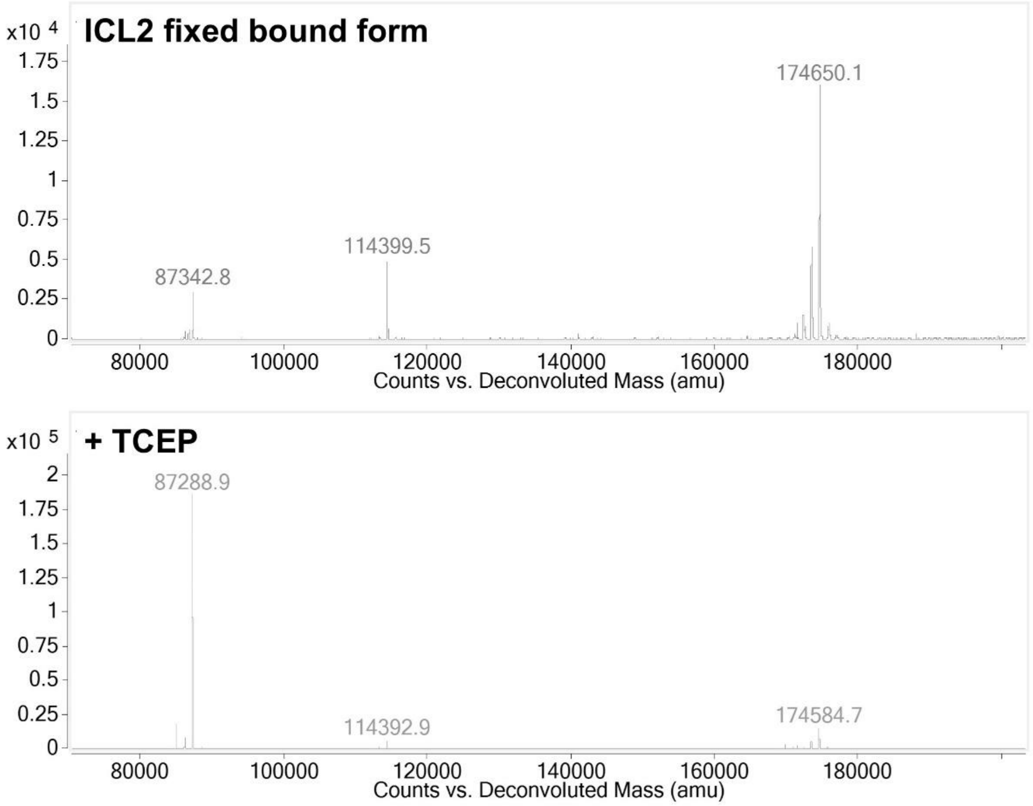


**Figure S9. Deconvolution spectra of the ICL2 fixed in the acetyl-CoA bound form (E640C/N731C/E733C) obtained via denaturing mass spectrometry.** The top spectrum represents the protein on its own, while the bottom spectrum represents the protein in the presence of TCEP. The average mass of the ICL2 fixed in the acetyl-CoA bound form is 87420 Da, while the observed mass was 87289 Da. The mass loss of 131 Da may correspond to the loss of the N-terminal methionine. Under denaturing conditions, the protein appears to form a covalent dimer (~170 kDa) that can be reduced to a monomer (~87 kDa) in the presence of TCEP. The calculated mass of the ICL2 fixed *apo* form dimer (no N-terminal methionine) with three disulfide bonds (-2 Da for each disulfide bond formation) is 174572 Da, and the observed mass is 174650 Da.

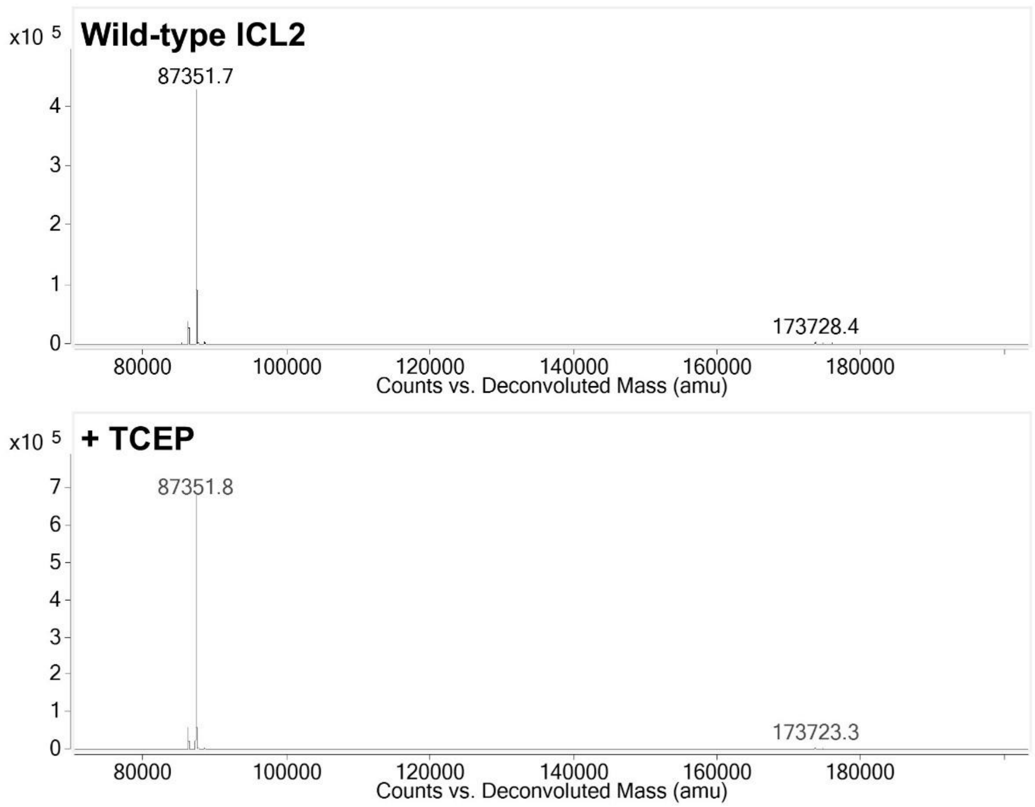


**Figure S10. Deconvolution spectra of wild-type ICL2 obtained via denaturing mass spectrometry.** The top spectrum represents the protein on its own, while the bottom spectrum represents the protein in the presence of TCEP. The average mass of the ICL2 fixed in the acetyl-CoA bound form is 87483 Da, while the observed mass is 87352 Da. The mass loss of 131 Da may correspond to the loss of the N-terminal methionine.

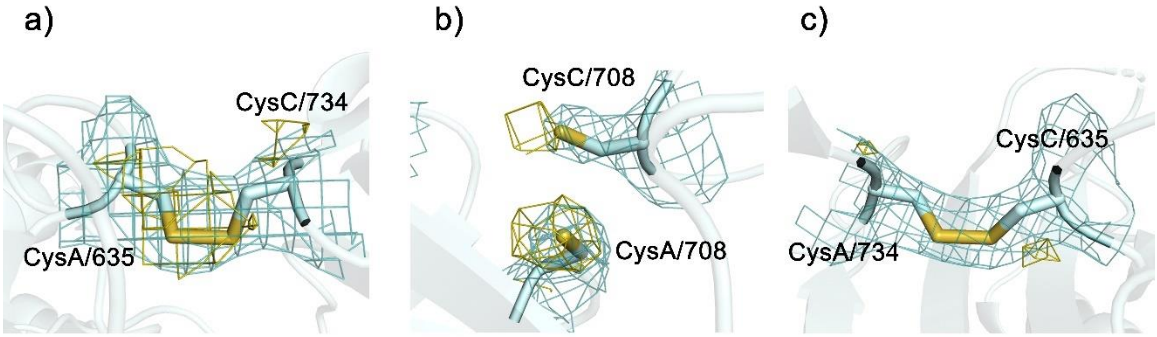


**Figure S11. Disulfide bonds formed in the ICL2 fixed *apo* form.** The mutated cysteines are highlighted in chains A and C of the crystal structure of the ICL2 fixed *apo* form (PDB: 9OBO). The 2Fo-Fc map is shown in teal, while the anomalous map is shown in yellow.

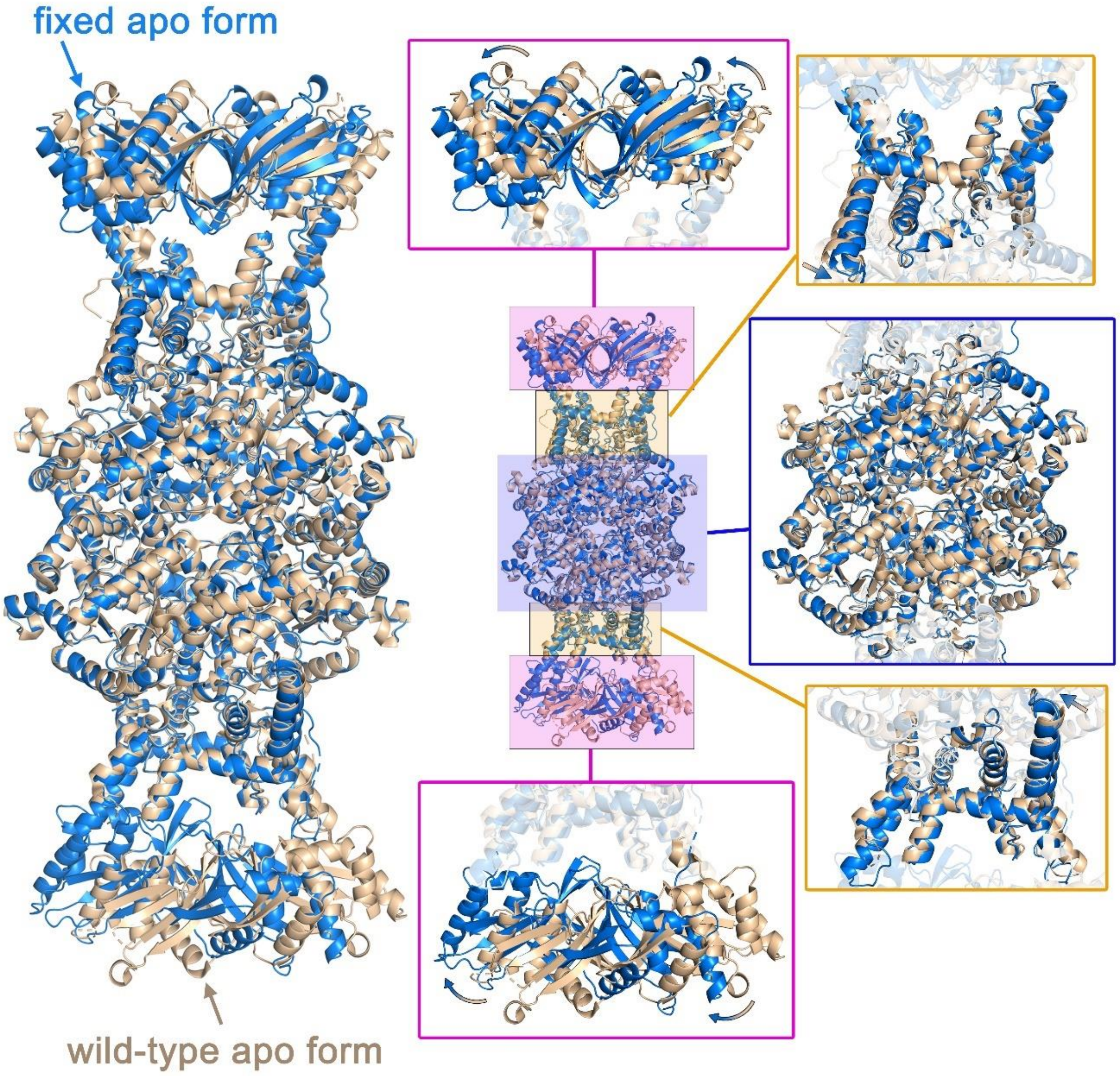


**Figure S12. Crystal structure of ICL2 fixed *apo* form vs wild-type ICL2.** The crystal structure of the ICL2 fixed *apo* form (blue; PDB: 9OBO) is overlaid with the crystal structure of *apo* ICL2 (wheat; PDB: 6EDW [17]). The different domains are zoomed in on the right-hand side with arrows indicating the direction of the shift in position. The C-terminal domains are highlighted in pink, the unique helical substructure in orange, and the N-terminal domains in blue.

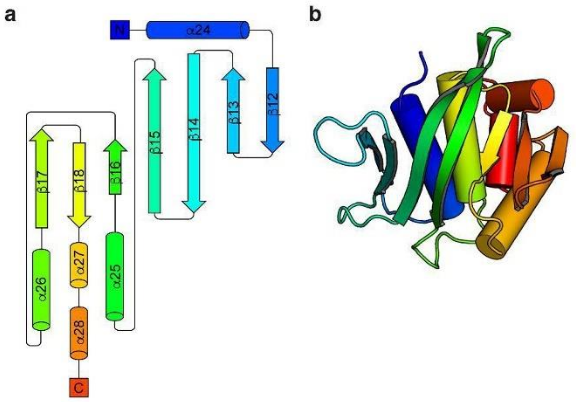


**Figure S13. Secondary structure of the ICL2 C-terminal domain (CTD). (**a) Topology diagram for the CTD. (b) Cartoon representation of the CTD coloured according to the topology diagram. These figures were taken from Bhusal *et al.* [17] as a reference for the numbering of secondary structures in the discussion of the manuscript.

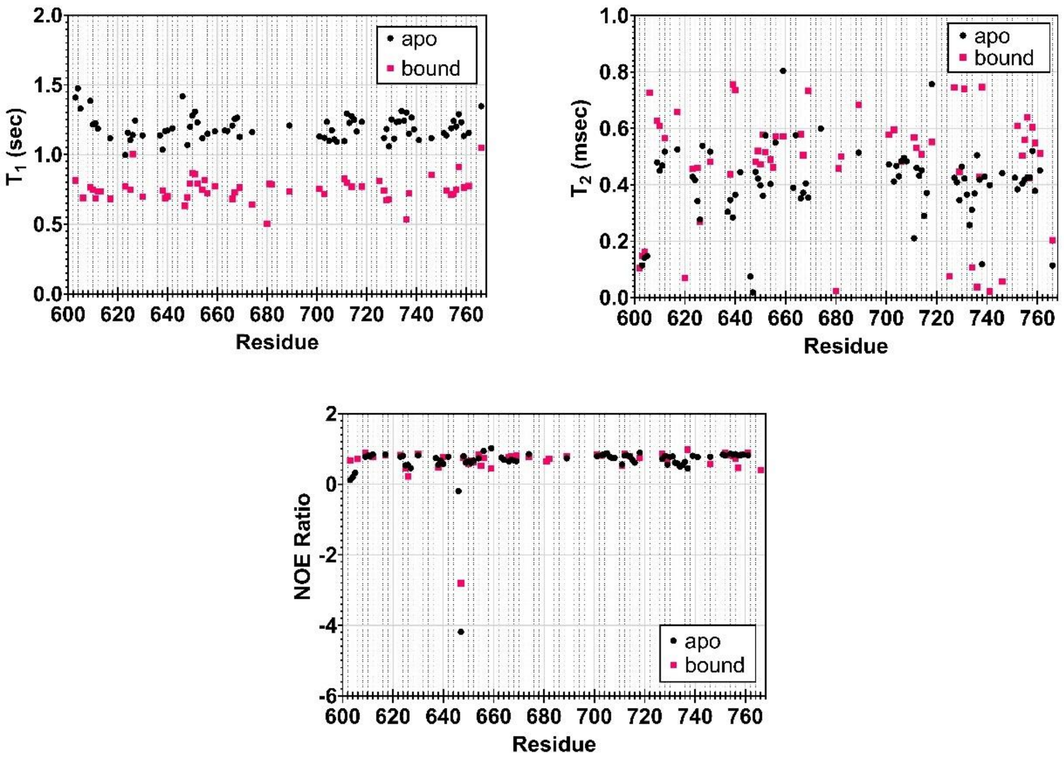


**Figure S14. $^{15}N$ relaxation experiments of *apo*- and acetyl-CoA bound isolated ICL2 C-terminal domain (CTD).** $^{15}N$ relaxation experiments shown in $T_1$ relaxation, $T_2$ relaxation, and $^{1}H$-$^{15}N$ NOE measurements of *apo* (black circles) and acetyl-CoA bound (pink squares) CTD. Experiments were conducted with 300 μM $^{2}H$, $^{15}N$-CTD with or without 2 mM acetyl-CoA. Samples were in 50 mM Tris-$d_{11}$ at pH 6.6 with 90% $H_2O$ and 10% $D_2O$ at 298 K.

a)

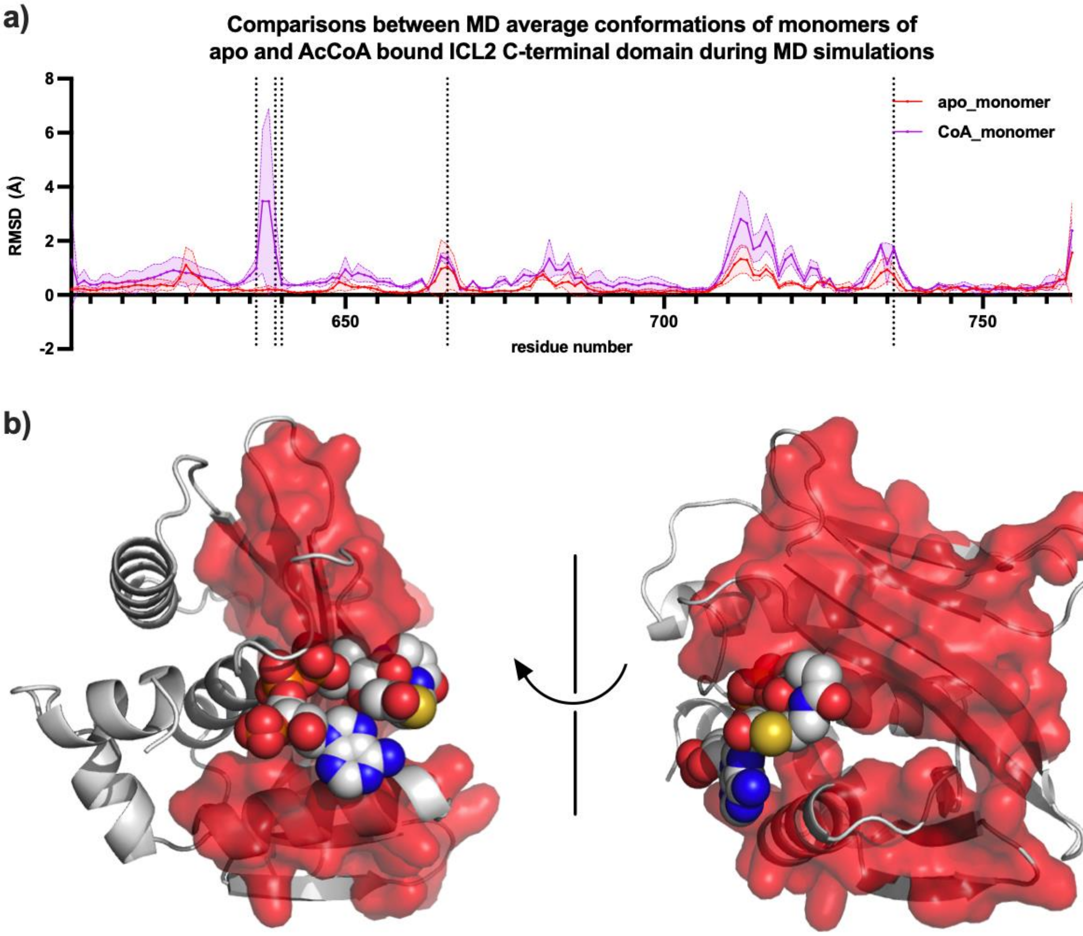


b)

**Figure S15. Conformational changes in the C-terminal domain (CTD) upon binding of acetyl-CoA.** The average structure of chain A in one of the MD simulations of the *apo*-ICL2 CTD was used as the reference structure for measurement of r.m.s.d. values. The average structures were all aligned to the reference structure using backbone atoms of residues 607 to 760. (a) r.m.s.d. values between the *apo* and acetyl-CoA-bound CTD. (b) The *apo* state CTD represented in cartoon form. The residues that showed the highest r.m.s.d. values between the monomer and dimer forms in the MD simulation are highlighted in red.

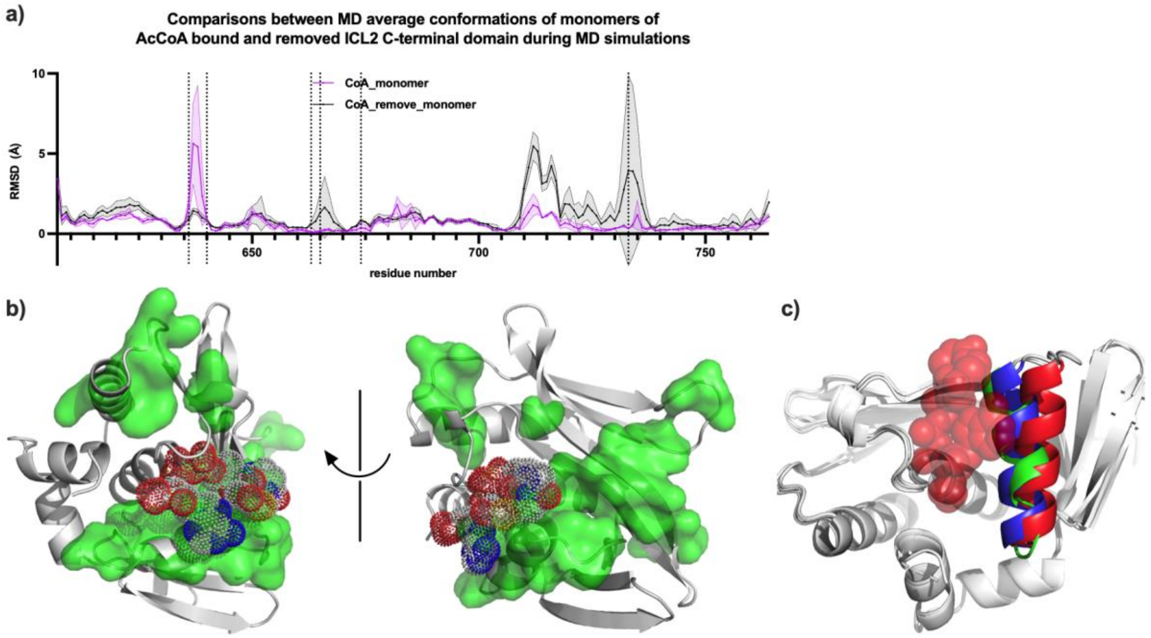


**Figure S16. Conformational changes in the C-terminal domain (CTD) upon binding of acetyl-CoA.** The average structure of chain A in the first MD simulation of the acetyl-CoA bound CTD was used as the reference structure for measurement of r.m.s.d. values. The average structures were all aligned to the reference structure using backbone atoms of residues 607 to 760. (a) r.m.s.d. values of the acetyl-CoA-bound CTD when acetyl-CoA was removed. (b) The bound state CTD represented in cartoon form. The residues that showed the highest r.m.s.d. values when acetyl-CoA was removed in the MD simulation are highlighted in green. Acetyl-CoA is illustrated as a ball model. (c) Conformations of α26 in the *apo* CTD (blue), acetyl-CoA-bound CTD (red) and acetyl-CoA-removed CTD (green). MD average structures from the first simulation of each system were superimposed. The protein is represented in cartoon form, while acetyl-CoA is represented by a ball model.

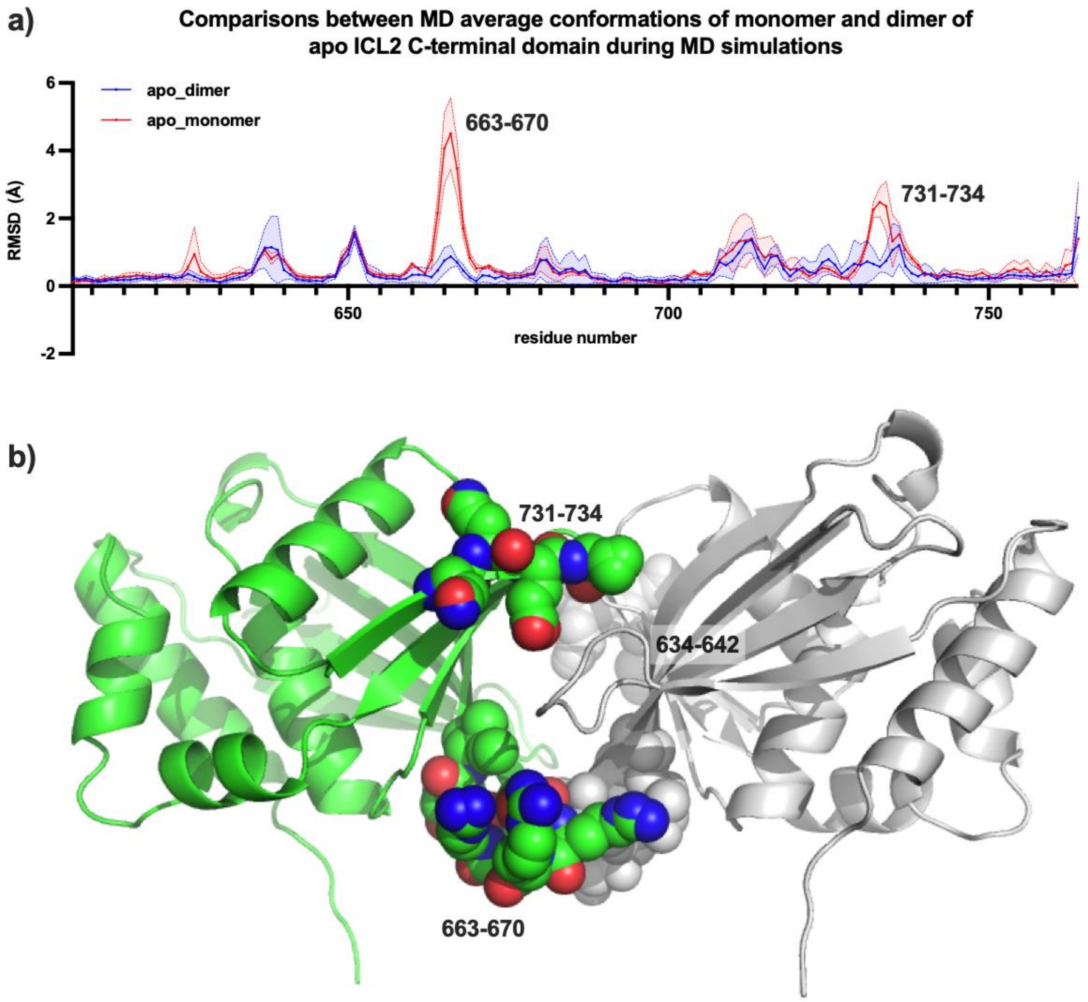


**Figure S17. Effect of dimerisation on local conformations in the *apo* state C-terminal domain (CTD) dimer.** The average structure of chain A in the first MD simulation of the *apo* state dimer was used as the reference structure for measurement of r.m.s.d. values. The average structures were all aligned to the reference structure using backbone atoms of residues 607 to 760. (a) r.m.s.d. values between the monomer and dimer forms of the *apo*-state CTD dimer. (b) The *apo* state CTD dimer represented in cartoon form. The residues that showed the highest r.m.s.d. values between the monomer and dimer forms in the MD simulation are highlighted as ball models.

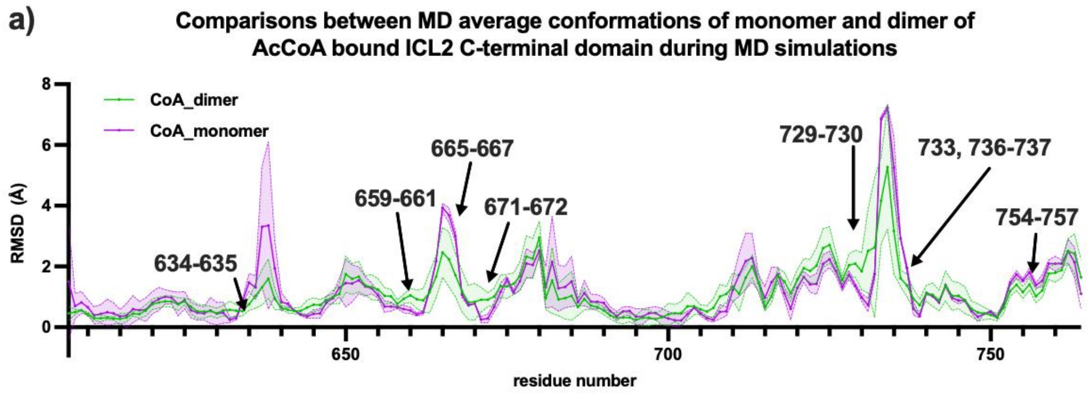


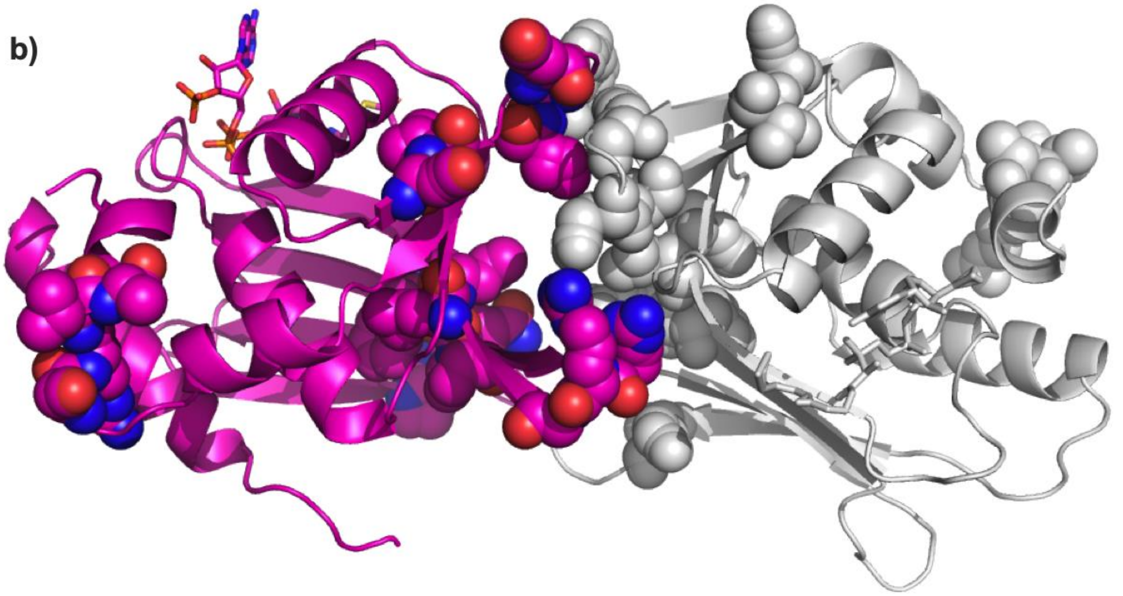


**Figure S18. Effect of dimerisation on local conformations in the presence of acetyl-CoA (AcCoA).** The average structure of chain A in the first MD simulation of AcCoA-bound dimer was used as the reference structure for measurement of r.m.s.d. values. The average structures were all aligned to the reference structure using backbone atoms of residues 607 to 760. (a) r.m.s.d. values between the monomer and dimer forms of the acetyl-CoA-bound C-terminal domain (CTD) dimer. (b) The bound state CTD dimer represented in cartoon form. The residues that showed the highest r.m.s.d. values between the monomer and dimer forms in the MD simulation are highlighted as ball models.

```
ICL2      1 MAIAETDTEVHTPFEQDFEKDVAATQRYFDSSRFAGIIRLYTARQ 45
ICL1                    1 MSVVGTPKSAEQIQQEWDTNPRWKDVTRTYSAED 35
ICL2ΔUHS    MAIAETDTEVHTPFEQDFEKDVAATQRYFDSSRFAGIIRLYTARQ

ICL2     46 VVEQRGTIPVDHIVAREAAGAFYERLRELFAARKSITTFGPYSPG 90
ICL1     36 VVALQGSVVEEHTLARRGAEVLWEQLHDL----EWVNALGALTGN 75
ICL2ΔUHS    VVEQRGTIPVDHIVAREAAGAFYERLRELFAARKSITTFGPYSPG

ICL2     91 QAVSMKRMGIEAIYLGGWATSAKGSSTEDPGPDLASYPLSQVPDD 135
ICL1     76 MAVQQVRAGLKAIYLSGWQVAGDANLSGHTYPDQSLYPANSVPQV 120
ICL2ΔUHS    QAVSMKRMGIEAIYLGGWATSAKGSSTEDPGPDLASYPLSQVPDD

ICL2    136 AAVLVRALLTADRNQHYLRLQMSERQRAATPAYDFRPFIIADADT 180
ICL1    121 VRRINNALQRADQ---------IAKIEGDTSVENWLAPIVADGEA 156
ICL2ΔUHS    AAVLVRALLTADRNQHYLRLQMSERQRAATPAYDFRPFIIADADT

ICL2    181 GHGGDPHVRNLIRRFVEVGVPGYHIEDQRPGTKKCGHQGGKVLVP 225
ICL1    157 GFGGALNVYELQKALIAAGVAGSHWEDQLASEKKCGHLGGKVLIP 201
ICL2ΔUHS    GHGGDPHVRNLIRRFVEVGVPGYHIEDQRPGTKKCGHQGGKVLVP

ICL2    226 SDEQIKRLNAARFQLDIMRVPGIIVARTDAEAANLIDSRADERDQ 270
ICL1    202 TQQHIRTLTSARLAADVADVPTVVIARTDAEAATLITSDVDERDQ 246
ICL2ΔUHS    SDEQIKRLNAARFQLDIMRVPGIIVARTDAEAANLIDSRADERDQ

ICL2    271 PFLLGATKLDVPSYKSCFLAMVRRFYELGVKELNGHLLYALGDSEY 315
ICL1    247 PFITGER 253 --------------------------------------
ICL2ΔUHS    PFITGER--------------------------------------

ICL2    316 AAAGGWLERQGIFGLVSDAVNAWREDGQQSIDGIFDQVESRFVA 360
ICL1        ---------------------------------------------
ICL2ΔUHS    ---------------------------------------------

ICL2    361 AWEDDAGLMTYGEAVADVLEFGQSEGEPIGMAPEEWRAFAARASL 405
ICL1        ---------------------------------------------
ICL2ΔUHS    ---------------------------------------------

ICL2    406 HAARAKAKELGADPPWDCELAKTPEGYYQIRGGIPYAIAKSLAAA 450
ICL1        ---------------------- 254 TREGFYRTKNGIEPCIARAKAYA 276
ICL2ΔUHS    ----------------------TPEGYYQIRGGIPYAIAKSLAAA

ICL2    451 PFADILWMETKTADLADARQFAEAIHAEFPDQMLAYNLSPSFNWD 495
ICL1    277 PFADLIWMETGTPDLEAARQFSEAVKAEYPDQMLAYNCSPSFNWK 321
ICL2ΔUHS    PFADILWMETKTADLADARQFAEAIHAEFPDQMLAYNLSPSFNWD

ICL2    496 TTGMTDEEMRRFPEELGKMGFVFNFITYGGHQIDGVAAEEFATAL 540
ICL1    322 KHLDDATIAKFQKELAAMGFKFQFITLAGFHALNYSMFDLAYGY 365
ICL2ΔUHS    TTGMTDEEMRRFPEELGKMGFVFNFITYGGHQIDGVAAEEFATAL

ICL2    541 RQDGMLALARLQRKMRLVESPYRTPQTLVGGPRSDAALAASSGRT 585
ICL1    366 AQNQMSAYVELQEREFAAEERGYTATKHQREVGAGYFDRIATTVD 410
ICL2ΔUHS    RQDGMLALARLQRKMRLVESPYRTPQTLVGGPRSDAALAASSGRT

ICL2    586 ATTKAMGKGSTQHQHLVQTEVPRKLLEEWLAMWSGHYQLKDKLRV 630
ICL1    411 PNSSTTALTGSTEEGQFH 428
ICL2ΔUHS    ATTKAMGKGSTQHQHLVQTEVPRKLLEEWLAMWSGHYQLKDKLRV

ICL2    631 QLRPQRAGSEVLELGIHGESDDKLANVIFQPIQDRRGRTILLVRD 675

ICL2ΔUHS    QLRPQRAGSEVLELGIHGESDDKLANVIFQPIQDRRGRTILLVRD

ICL2    676 QNTFGAELRQKRLMTLIHLWLVHRFKAQAVHYVTPTDDNLYQTSK 720

ICL2ΔUHS    QNTFGAELRQKRLMTLIHLWLVHRFKAQAVHYVTPTDDNLYQTSK

ICL2    721 MKSHGIFTEVNQEVGEIIVAEVNHPRIAELLTPDRVALRKLITKEA 766

ICL2ΔUHS    MKSHGIFTEVNQEVGEIIVAEVNHPRIAELLTPDRVALRKLITKEA
```

**Figure S19. Sequence of ICL2 with the unique helical substructure deleted (ICL2ΔUHS) compared with ICL1 and ICL2.** The sequence of full-length *Mycobacterium tuberculosis* CDC1551 ICL2 is in black. The sequence of full-length *M. tuberculosis* H37Rv ICL1 is in orange. The sequence of ICL2ΔUHS is in blue. The region near the deletion was mutated to mimic the *M. tuberculosis* ICL1 sequence to promote a more likely formation of a folded protein. This region is highlighted in pink. The sequence alignment of ICL1 and ICL2 is based on a BLAST (https://blast.ncbi.nlm.nih.gov/BlastAlign.cgi) alignment of the two proteins.

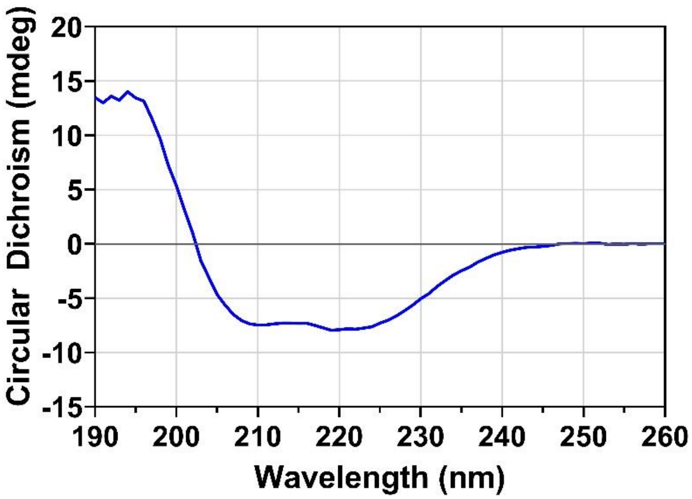


**Figure S20. Circular dichroism (CD) spectra of ICL2 with the unique helical substructure deleted (ICL2ΔUHS).** 0.1 mg/mL protein in 10 mM sodium phosphate (pH 7.5) was used. Measurements were performed at 25 °C.

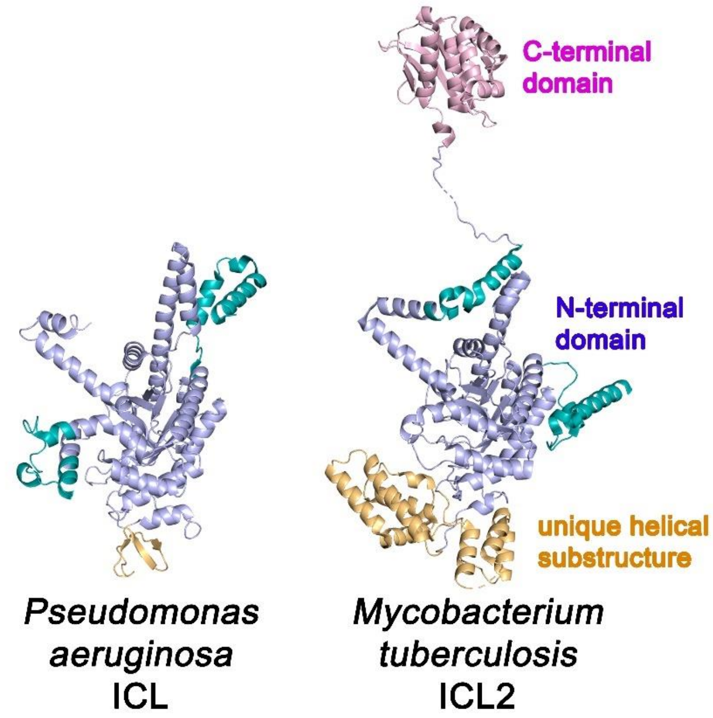


**Figure S21. Comparison of *Pseudomonas aeruginosa* ICL with *M. tuberculosis* ICL2.** One monomeric subunit from *P. aeruginosa* ICL (PDB: 6G1O [18]) was compared with one monomeric subunit from *M. tuberculosis* ICL2 (PDB: 6EDW [17]). The protein structures are coloured according to the protein features/domains. Portions in the N-terminal domain that did not match between the two structures are coloured in teal.

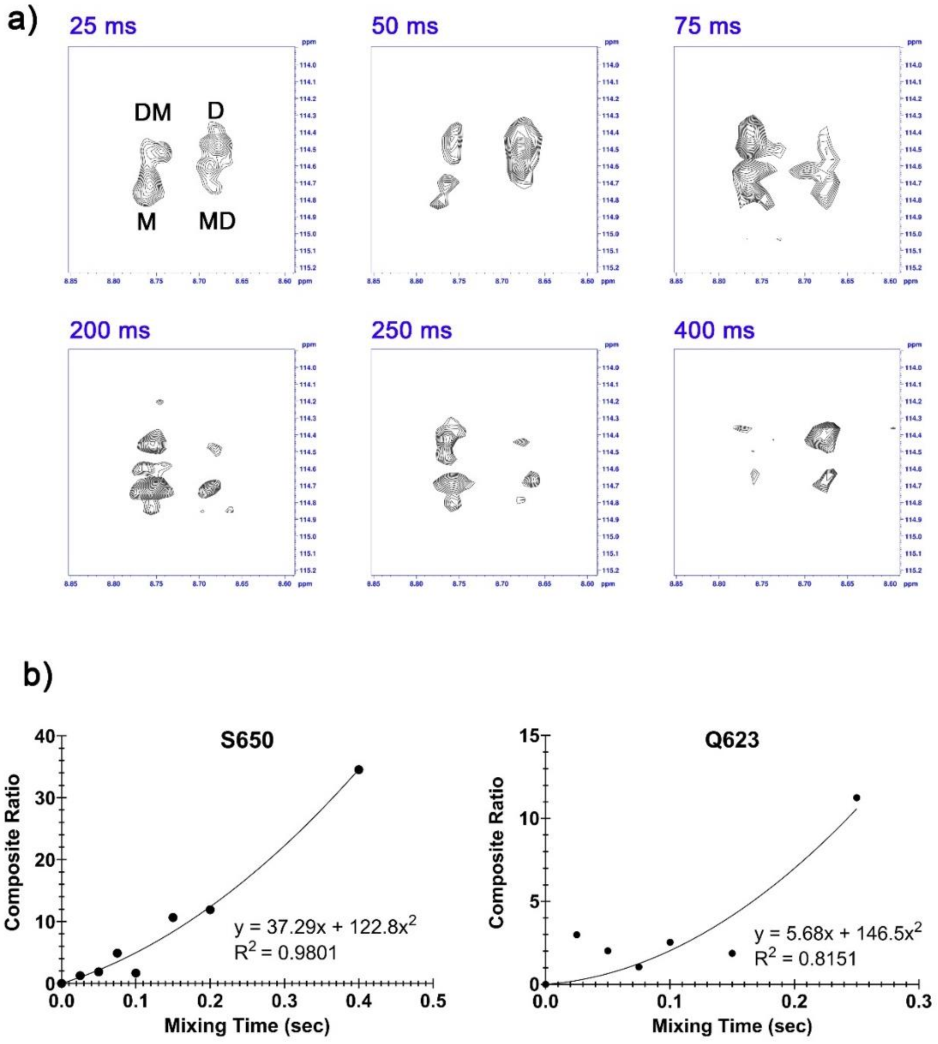


**Figure S22. ZZ-exchange spectra of ICL2 C-terminal domain (CTD) in the presence of acetyl-CoA. (**a) A series of ZZ-exchange spectra of S650 over different mixing times. The experiment was conducted with 300 μM of $^2H$, $^{15}N$-CTD and 500 μM acetyl-CoA in 50 mM Tris-$d_{11}$ (pH 6.6) with 90% $H_2O$ / 10% $D_2O$ at 298 K. M represents the monomer peak, D the dimer peak, MD the monomer to dimer interconversion, and DM the dimer to monomer interconversion. (b) The composite peak intensity ratio at different mixing times for S650 and Q623. The data was fitted to a quadratic equation using Graphpad Prism 8.0.

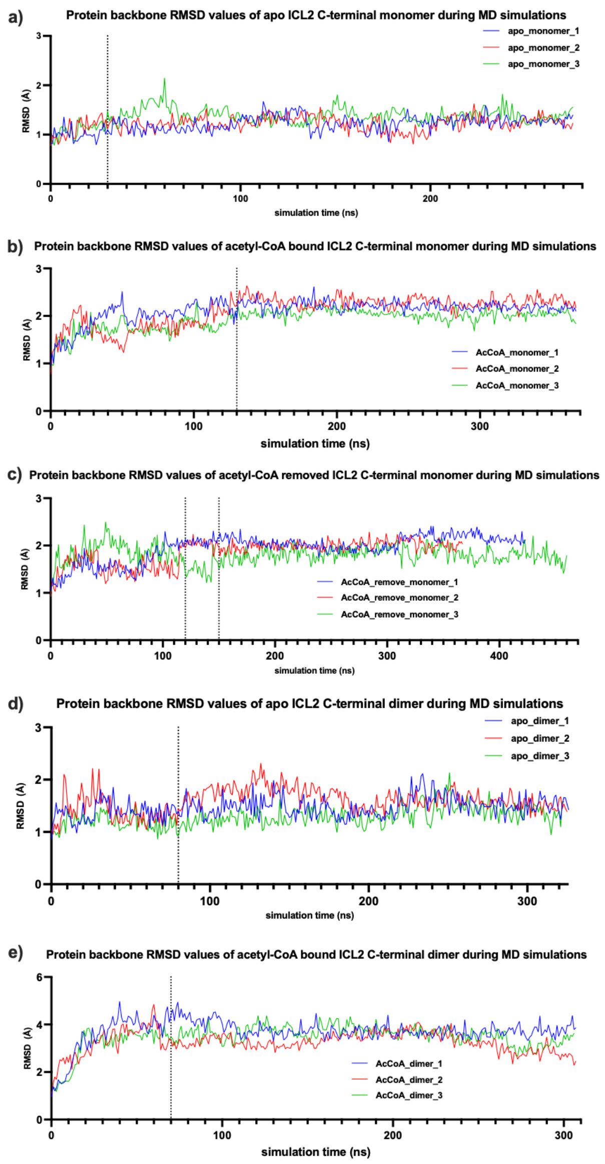


**Figure S23. Protein backbone r.m.s.d. values during MD simulations.** (a) C-terminal domain (CTD) monomer in *apo* state. Simulations equilibrated after 30 ns. Total trajectory time used for analysis was 737.6 ns. (b) CTD monomer with acetyl-CoA bound. Simulations equilibrated after 130 ns. Total trajectory time used for analysis was 712.3 ns. (c) CTD monomer starting from acetyl-CoA bound conformation but with acetyl-CoA removed. Simulations 1 and 2 equilibrated after 120 ns. Simulation 3 equilibrated after 150 ns. Total trajectory time used was 861.7 ns. (d) CTD dimer in the *apo* state. Simulations equilibrated after 80 ns. Total trajectory time used was 736 ns. (e) CTD dimer with acetyl-CoA bound. Simulations equilibrated after 70 ns. Total trajectory time used was 711.3 ns.

**Table S1. Interface hydrogen bonds formed during MD simulations of different states of C-terminal domain (CTD) dimers.** Hydrogen bonds with average occupancies above 30% are shown. The interaction of R665 is highlighted in pink. R665 interacts with the neighbouring CTD in the *apo* state but interacts with acetyl-CoA in the acetyl-CoA-bound state (Table S2).

| Donor Residue | Acceptor Residue | *apo* State Dimer | AcCoA State Dimer |
|---|---|---|---|
| A: R636 (Side Chain) | B: E733 (Side Chain) | 64.53% | |
| B: R636 (Side Chain) | A: E733 (Side Chain) | 52.23% | |
| A: R636 (Side Chain) | B: E640 (Side Chain) | | 45.70% |
| B: R636 (Side Chain) | A: E640 (Side Chain) | | 34.89% |
| **B: R665 (Side Chain)** | **A: E640 (Side Chain)** | **67.36%** | |
| **A: R665 (Side Chain)** | **B: E640 (Side Chain)** | **56.78%** | |
| A: R666 (Side Chain) | B: E640 (Side Chain) | 46.59% | |
| B: R666 (Side Chain) | A: E640 (Side Chain) | 35.08% | |
| A: R666 (Side Chain) | B: E736 (Side Chain) | | 36.98% |
| B: R666 (Side Chain) | A: E736 (Side Chain) | | 50.59% |
| B: R668 (Side Chain) | A: E736 (Side Chain) | | |
| B: R674 (Side Chain) | A: E733 (Side Chain) | 46.28% | |
| A: R674 (Side Chain) | B: E733 (Side Chain) | 37.16% | |
| B: N663 (Main Chain) | A: N663 (Main Chain) | 43.73% | |
| A: S639 (Side Chain) | B: E640 (Side Chain) | | 32.10% |
| B: S639 (Side Chain) | A: E640 (Side Chain) | | 39.09% |

**Table S2. Hydrogen bonds formed between acetyl-CoA and the C-terminal domain during MD simulations.** Hydrogen bonds with average occupancies above 20% are shown. The interaction of R665 with the acetyl group in acetyl-CoA is highlighted in pink.

| donor | acceptor | CoA_dimer_A | CoA_dimer_C | CoA_monomer |
|---|---|---|---|---|
| R687-Side | COA1-Side | 84.72% | 90.61% | 84.03% |
| K720-Side | COA1-Side | 61.42% | 63.82% | 61.91% |
| K764-Side | COA1-Side | 54.43% | 57.79% | 48.46% |
| R684-Side | COA1-Side | 53.89% | 47.32% | 72.41% |
| Q685-Main | COA1-Side | 41.34% | 54.68% | 51.96% |
| T678-Side | COA1-Side | 39.61% | 30.74% | 27.59% |
| T690-Side | COA1-Side | 34.28% | 24.23% | 51.68% |
| **R665-Side** | **COA1-Side** | **25.56%** | **27.36%** | |
| Q717-Side | COA1-Side | | | 21.85% |
| M689-Main | COA1-Side | | 22.56% | |

**Table S3. Data collection and refinement statistics for the ICL2 fixed *apo* form.**

| | ICL2 fixed *apo* form | Anomalous scattering |
|---|---|---|
| **Data Collection** | | |
| Wavelength (Å) | 0.954 | 1.459 |
| Temperature (K) | 100 | |
| Diffraction Source | Australian Synchrotron MX2 | |
| Detector | Eiger 16M | |
| Crystal-to-detector distance (mm) | 379 | |
| Space Group | $P\,2_1$ | $P\,2_1$ |
| *a*, *b*, *c* (Å) | 103.70, 170.86, 104.37 | 103.68, 171.37, 104.78 |
| *α*, *β*, *γ* (°) | 90.00, 94.14, 90.00 | 90.00, 94.12, 90.00 |
| Resolution (Å) | 49.50 – 2.60 (2.64 – 2.60) | 50.12 – 3.50 (3.62 – 3.50) |
| Total no. of reflections | 489810 (25366) | 927970 (64884) |
| No. of unique reflections | 110858 (5467) | 45313 (3639) |
| Completeness (%) | 99.8 (100.0) | 98.0 (80.4) |
| Redundancy | 4.4 (4.6) | 20.5 (17.8) |
| Average $I/\sigma I$ | 10.3 (2.2) | 17.7 (6.0) |
| $R_{merge}$ (%) | 7.6 (67.9) | 12.6 (35.7) |
| $R_{pim}$ (%) | 4.1 (35.4) | 2.9 (8.4) |
| $CC_{1/2}$ | 0.995 (0.766) | 0.999 (0.980) |
| **Refinement** | | |
| Resolution (Å) | 49.55 – 2.60 (2.67 – 2.60) | |
| $R_{work}$ (%) | 17.5 | |
| $R_{free}$ (%) | 22.4 | |
| No of reflections in working set | 105196 | |
| No. of reflections in test set | 5620 (5.1%) | |
| No. of protein molecules per asu | 4 | |
| No. of other groups per asu | 1 (succinate) | |
| R.m.s.d. bond lengths (Å) | 0.003 | |
| R.m.s.d. bond angles (°) | 0.873 | |
| *Average B-factors (Å²)*[b] | | |
| Protein | 75.6 | |
| Water | 56.5 | |
| Other | 66.5 | |
| *Ramachandran Plot*[c] | | |
| Most favoured regions (%) | 97.1 | |
| Additional allowed regions (%) | 2.7 | |
| Disallowed regions (%) | 0.2 | |
| PDB code | 9OBO | |

[a] Values in brackets are for the highest-resolution shell. [b] Calculated using $B_{average}$ from the CCP4 Suite. [c] Calculated using MolProbity.

**Table S4. Summary of backbone amide assignment of $^{13}C$, $^{15}N$-CTD.** Chemical shifts are given in ppm. The gaps indicate missing assignments. The N-terminal polyhistidine tag is omitted. Among the 138 peaks picked from the $^1H$-$^{15}N$ HSQC, 97 were assigned to the 158 non-proline or non-N-terminal amino acids.

| Residue | $^1H$ | $^{15}N$ | CO | Cα | Cβ |
|---|---|---|---|---|---|
| L 601 | 8.21 | 123.71 | 177.04 | 55.23 | 42.28 |
| V 602 | 7.97 | 121.24 | 175.96 | 62.33 | 32.58 |
| Q 603 | 8.4 | 124.32 | | 55.79 | 29.42 |
| T 604 | 8.07 | 115.73 | 174.23 | 61.94 | 69.68 |
| E 605 | 8.26 | 123.57 | 176.12 | 56.28 | 30.47 |
| V 606 | 8.13 | 125.22 | 173.45 | 60.40 | 31.87 |
| P 607 | | | | | |
| R 608 | | | | | |
| K 609 | 8.14 | 117.47 | 176.62 | 57.89 | 32.32 |
| L 610 | 7 | 118.70 | | 42.93 | |
| L 611 | 5.75 | 114.46 | 178.53 | 56.30 | 37.99 |
| E 612 | 7.28 | 116.91 | 179.62 | 60.72 | 28.57 |
| E 613 | 8.34 | 121.69 | 180.72 | 59.94 | |
| W 614 | | | | | |
| L 615 | 9.05 | 117.72 | 178.89 | 57.33 | 40.61 |
| A 616 | | | | | |
| M 617 | 7.77 | 120.47 | 178.89 | 59.38 | 33.97 |
| W 618 | | | | | |
| S 619 | | | | | |
| G 620 | | | | | |
| H 621 | | | | | |
| Y 622 | | | | | |
| Q 623 | 7.48 | 115.07 | 175.09 | 56.69 | 25.4 |
| L 624 | 8.08 | 119.10 | 178.19 | 54.32 | 42.53 |
| K 625 | 8.56 | 121.84 | 176.57 | 56.96 | 32.61 |
| D 626 | 7.59 | 119.31 | 175.58 | 54.17 | 42.2 |
| K 627 | 8.94 | 124.37 | | 55.98 | 32.26 |
| L 628 | | | | | |
| R 629 | 9.14 | 122.55 | 174.78 | 53.94 | 33.59 |
| V 630 | 9.33 | 121.58 | 175.18 | 60.81 | |
| Q 631 | | | | | |
| L 632 | | | | | |
| R 633 | | | | | |
| P 634 | | | | | |
| Q 635 | 9.12 | 124.39 | 177.81 | 59.18 | 28.96 |
| R 636 | 7.54 | 111.24 | 174.59 | | 32.85 |
| A 637 | 8.51 | 124.13 | 179.1 | 54.07 | 17.65 |
| G 638 | 8.82 | 111.16 | | 45.40 | |

| | | | | | |
|---|---|---|---|---|---|
| S 639 | 7.78 | 113.63 | 173.93 | 57.54 | 64.84 |
| E 640 | 8.8 | 123.67 | 175.77 | 56.85 | 30.13 |
| V 641 | | | | | |
| L 642 | 8.6 | 127.72 | 174.35 | 53.42 | 44.50 |
| E 643 | 8.74 | 118.01 | 175.86 | 54.56 | 32.99 |
| L 644 | 9.47 | 131.82 | 175.48 | 53.22 | 43.91 |
| G 645 | | | | | |
| I 646 | | | | | |
| H 647 | | | | | |
| G 648 | 8.33 | 108.44 | 175.95 | 43.74 | |
| E 649 | 9.67 | 124.11 | 178.15 | 59.45 | 29.76 |
| S 650 | 8.69 | 114.59 | 174.26 | 58.36 | 63.32 |
| D 651 | 8.31 | 116.36 | 176.15 | 56.59 | 40.22 |
| D 652 | 8.29 | 120.84 | 175.27 | 54.98 | |
| K 653 | | | | | |
| L 654 | | | | | |
| A 655 | | | | | |
| N 656 | 9.36 | 118.03 | 172.83 | 52.57 | 40.98 |
| V 657 | | | | | |
| I 658 | 9.21 | 128.92 | 176.31 | 60.1 | 38.36 |
| F 659 | 9.46 | 125.64 | 170.62 | 55.77 | 43.19 |
| Q 660 | 8.89 | 118.07 | 173.23 | 52.31 | 33.42 |
| P 661 | | | | | |
| I 662 | 8.87 | 122.25 | 174.06 | 60.07 | 42.32 |
| Q 663 | 7.97 | 118.77 | 176.73 | 54.09 | 31.73 |
| D 664 | 8.68 | 122.68 | 178.55 | 52.17 | 41.24 |
| R 665 | 8.35 | 116.40 | 177.31 | 58.28 | 29.54 |
| R 666 | 7.92 | 118.21 | 176.37 | 55.60 | 30.71 |
| G 667 | 8.07 | 108.53 | 174.25 | 46.85 | |
| R 668 | 8.47 | 121.06 | 176.29 | 55.38 | 30.92 |
| T 669 | 8.9 | 119.51 | 173.51 | 64.02 | 68.87 |
| I 670 | 9.16 | 125.55 | 174.77 | 58.24 | 41.81 |
| L 671 | | | | | |
| L 672 | 8.61 | 127.02 | 175.71 | 53.83 | 42.42 |
| V 673 | 9.58 | 128.35 | 176.07 | 63.33 | 31.49 |
| R 674 | 9.08 | 128.80 | | 57.20 | 31.20 |
| D 675 | | | | | |
| Q 676 | | | | | |
| N 677 | | | | | |
| T 678 | | | | | |
| F 679 | | | | | |
| G 680 | | | | | |
| A 681 | | | | | |

| | | | | | |
|---|---|---|---|---|---|
| E 682 | | | | | |
| L 683 | | | | | |
| R 684 | | | | | |
| Q 685 | | | | | |
| K 686 | | | | | |
| R 687 | | | | | |
| L 688 | | | | | |
| M 689 | 8.2 | 123.30 | | 58.07 | 28.79 |
| T 690 | | | | | |
| L 691 | | | | | |
| I 692 | | | | | |
| H 693 | | | | | |
| L 694 | | | | | |
| W 695 | | | | | |
| L 696 | | | | | |
| V 697 | 8.78 | 115.60 | 178.27 | 66.52 | |
| H 698 | 7.59 | 120.46 | 176.88 | 57.87 | 31.42 |
| R 699 | 8.88 | 119.34 | 177.64 | | 27.87 |
| F 700 | | | | | |
| K 701 | 7.41 | 118.68 | 173.29 | 55.87 | 28.47 |
| A 702 | 7.7 | 118.00 | 177.6 | 51.97 | 20.08 |
| Q 703 | 9.18 | 117.22 | 175.39 | 56.14 | 29.73 |
| A 704 | 7.64 | 116.89 | 174.75 | 51.20 | 22.49 |
| V 705 | 9.28 | 120.51 | 173.62 | 61.49 | 34.16 |
| H 706 | 9.31 | 127.06 | 174.19 | 52.53 | |
| Y 707 | 9.52 | 121.86 | 176.14 | 57.19 | 38.87 |
| V 708 | 7.38 | 117.59 | 174.69 | 64.5 | 32.07 |
| T 709 | | | | | |
| P 710 | | | | | |
| T 711 | 7.72 | 112.99 | 175.33 | 61.93 | 70.98 |
| D 712 | 8.88 | 121.27 | 179.47 | 57.96 | 39.84 |
| D 713 | 8.72 | 118.29 | 178.16 | 57.31 | 41.79 |
| N 714 | 8.16 | 119.59 | 179.16 | 55.95 | 36.59 |
| L 715 | 8.32 | 123.99 | 178.29 | 58.28 | 40.93 |
| Y 716 | 8.13 | 121.04 | | 61.60 | 38.30 |
| Q 717 | | | | | |
| T 718 | 7.86 | 109.29 | | 66.53 | 68.39 |
| S 719 | | | | | |
| K 720 | | | | | |
| M 721 | | | | | |
| K 722 | | | | | |
| S 723 | | | | | |
| H 724 | | | | | |

| | | | | | |
|---|---|---|---|---|---|
| G 725 | 8.35 | 110.10 | | 45.19 | |
| I 726 | | | | | |
| F 727 | 6.39 | 111.75 | 176.47 | 52.65 | 41.93 |
| T 728 | 8.75 | 113.09 | 174.62 | 62.81 | 68.29 |
| E 729 | 7.08 | 116.74 | 174.05 | 54.80 | 33.11 |
| V 730 | 8.59 | 122.67 | 174.42 | 61.70 | 34.02 |
| N 731 | 8.84 | 124.58 | 173.44 | 52.26 | 42.54 |
| Q 732 | 8.65 | 121.59 | 175.43 | 30.16 | |
| E 733 | 8.54 | 125.23 | 176.35 | 56.33 | 31.71 |
| V 734 | 8.21 | 120.69 | | 63.96 | |
| G 735 | 8.64 | 113.23 | 174.22 | 45.30 | |
| E 736 | 8.23 | 115.98 | 174.35 | 57.39 | 28.30 |
| I 737 | 6.85 | 115.40 | 175.74 | 39.73 | |
| I 738 | 8.05 | 123.88 | 174.85 | 60.56 | 39.34 |
| V 739 | 8.76 | 127.22 | 173.92 | 62.09 | 33.26 |
| A 740 | 9.13 | 132.76 | 175.47 | 49.65 | 19.11 |
| E 741 | 8.05 | 123.88 | 176.86 | 56.00 | 30.76 |
| V 742 | | | | | |
| N 743 | | | | | |
| H 744 | | | | | |
| P 745 | | | | | |
| R 746 | 8.38 | 121.75 | 176.83 | 56.37 | 31.06 |
| I 747 | | | | | |
| A 748 | | | | | |
| E 749 | | | | | |
| L 750 | | | | | |
| L 751 | 7.70 | 119.32 | 176.76 | 54.40 | 41.29 |
| T 752 | 7.33 | 119.09 | 176.05 | 62.94 | 68.99 |
| P 753 | | | | | |
| D 754 | 7.15 | 114.96 | 176.61 | 55.27 | 39.84 |
| R 755 | 8.05 | 112.83 | 175.89 | 58.55 | 27.72 |
| V 756 | 8.11 | 123.13 | 178.73 | 67.71 | 31.19 |
| A 757 | 9.16 | 123.63 | 181.11 | 54.84 | 17.18 |
| L 758 | 8.83 | 122.74 | 177.91 | 57.78 | 40.92 |
| R 759 | 8.42 | 119.05 | 179.26 | 60.56 | 29.50 |
| K 760 | | | | | |
| L 761 | 7.42 | 120.12 | 178.47 | 58.01 | 41.27 |
| I 762 | | | | | |
| T 763 | | | | | |
| K 764 | | | | | |
| E 765 | | | | | |
| A 766 | | | | | |

**Table S5. Transfer of $^{1}$H and $^{15}$N assignments from *apo* $^{2}$H, $^{15}$N-CTD to acetyl-CoA bound $^{2}$H, $^{15}$N-CTD.** Chemical shifts are given in ppm. Gaps indicate missing assignments or transfers. Nine weak resonances could not be transferred, and 27 peaks in the acetyl-C-A-bound $^{1}$H-$^{15}$N HSQC spectrum could not be assigned.

| | ***Apo* $^{2}$H, $^{15}$N-CTD** | | **Acetyl-CoA-bound $^{2}$H, $^{15}$N-CTD** | |
|---|---|---|---|---|
| **Residue** | **$^{1}$H** | **$^{15}$N** | **$^{1}$H** | **$^{15}$N** |
| L 601 | 8.19 | 123.48 | 8.20 | 123.90 |
| V 602 | 7.96 | 121.07 | 7.94 | 120.83 |
| Q 603 | 8.35 | 124.38 | 8.40 | 124.19 |
| T 604 | 8.06 | 115.72 | | |
| E 605 | 8.26 | 123.58 | 8.28 | 123.33 |
| V 606 | 8.13 | 125.16 | 8.00 | 125.21 |
| P 607 | | | | |
| R 608 | | | | |
| K 609 | 8.14 | 117.46 | 8.07 | 117.39 |
| L 610 | 7.00 | 118.63 | 7.01 | 118.79 |
| L 611 | 5.75 | 114.39 | 5.74 | 114.61 |
| E 612 | 7.26 | 116.81 | 7.36 | 116.88 |
| E 613 | 8.35 | 121.79 | 8.33 | 121.80 |
| W 614 | | | | |
| L 615 | 9.05 | 117.76 | | |
| A 616 | | | | |
| M 617 | 7.76 | 120.44 | 7.79 | 120.47 |
| W 618 | | | | |
| S 619 | | | | |
| G 620 | | | | |
| H 621 | | | | |
| Y 622 | | | | |
| Q 623 | 7.49 | 115.26 | 7.40 | 114.87 |
| L 624 | 8.11 | 119.11 | 8.06 | 118.44 |
| K 625 | 8.56 | 121.61 | 8.31 | 122.13 |
| D 626 | 7.61 | 119.21 | 7.58 | 119.51 |
| K 627 | 8.96 | 124.44 | 8.62 | 124.24 |
| L 628 | | | | |
| R 629 | 9.15 | 122.45 | 9.16 | 122.63 |
| V 630 | 9.36 | 121.59 | 9.30 | 121.36 |
| Q 631 | | | | |
| L 632 | | | | |
| R 633 | | | | |
| P 634 | | | | |
| Q 635 | 9.13 | 124.50 | 9.09 | 123.89 |
| R 636 | 7.53 | 111.22 | 7.56 | 111.24 |
| A 637 | 8.44 | 123.87 | 8.52 | 122.87 |

| | | | | |
|---|---|---|---|---|
| G 638 | 8.81 | 111.11 | | |
| S 639 | 7.78 | 113.64 | 7.79 | 113.68 |
| E 640 | 8.80 | 123.71 | 8.80 | 123.60 |
| V 641 | | | | |
| L 642 | 8.59 | 127.67 | 8.95 | 127.65 |
| E 643 | 8.74 | 118.04 | 8.73 | 117.77 |
| L 644 | 9.47 | 131.60 | | |
| G 645 | | | | |
| I 646 | | | | |
| H 647 | | | | |
| G 648 | 8.30 | 108.21 | 8.30 | 108.31 |
| E 649 | 9.66 | 123.89 | 9.82 | 124.10 |
| S 650 | 8.69 | 114.43 | 8.77 | 114.80 |
| D 651 | 8.30 | 116.36 | 8.31 | 116.24 |
| D 652 | 8.31 | 120.92 | 8.28 | 120.88 |
| K 653 | | | | |
| L 654 | | | | |
| A 655 | | | | |
| N 656 | 9.36 | 117.88 | 9.50 | 118.04 |
| V 657 | | | | |
| I 658 | 9.21 | 128.89 | | |
| F 659 | 9.46 | 125.55 | | |
| Q 660 | 8.89 | 118.02 | | |
| P 661 | | | | |
| I 662 | 8.87 | 122.28 | 8.75 | 122.26 |
| Q 663 | 7.96 | 118.71 | 7.97 | 118.78 |
| D 664 | 8.68 | 122.62 | 8.69 | 122.62 |
| R 665 | 8.33 | 116.47 | 8.31 | 117.13 |
| R 666 | 7.92 | 118.10 | 7.91 | 118.09 |
| G 667 | 8.07 | 108.52 | 8.06 | 108.5 |
| R 668 | 8.50 | 121.04 | 8.47 | 121.00 |
| T 669 | 8.89 | 119.49 | 8.89 | 119.52 |
| I 670 | 9.17 | 125.48 | 9.16 | 124.76 |
| L 671 | | | | |
| L 672 | 8.69 | 126.93 | 8.63 | 125.73 |
| V 673 | 9.58 | 128.34 | 9.58 | 128.34 |
| R 674 | 9.06 | 128.78 | 9.07 | 127.79 |
| D 675 | | | | |
| Q 676 | | | | |
| N 677 | | | | |
| T 678 | | | | |
| F 679 | | | | |
| G 680 | | | | |

| | | | | |
|---|---|---|---|---|
| A 681 | | | | |
| E 682 | | | | |
| L 683 | | | | |
| R 684 | | | | |
| Q 685 | | | | |
| K 686 | | | | |
| R 687 | | | | |
| L 688 | | | | |
| M 689 | 8.19 | 123.09 | 8.20 | 123.65 |
| T 690 | | | | |
| L 691 | | | | |
| I 692 | | | | |
| H 693 | | | | |
| L 694 | | | | |
| W 695 | | | | |
| L 696 | | | | |
| V 697 | 8.78 | 115.59 | | |
| H 698 | 7.58 | 120.47 | | |
| R 699 | 8.84 | 119.20 | 8.89 | 118.14 |
| F 700 | | | | |
| K 701 | 7.41 | 118.60 | 7.42 | 118.57 |
| A 702 | 7.69 | 118.01 | 7.72 | 118.03 |
| Q 703 | 9.19 | 117.05 | 9.17 | 117.25 |
| A 704 | 7.66 | 116.79 | 7.64 | 116.84 |
| V 705 | 9.30 | 120.48 | 9.25 | 120.50 |
| H 706 | 9.32 | 126.92 | 9.30 | 127.29 |
| Y 707 | 9.52 | 121.82 | 9.37 | 121.78 |
| V 708 | 7.38 | 117.64 | 7.49 | 117.40 |
| T 709 | | | | |
| P 710 | | | | |
| T 711 | 7.73 | 112.79 | 7.62 | 114.90 |
| D 712 | 8.88 | 121.24 | 8.87 | 121.19 |
| D 713 | 8.72 | 118.18 | 8.72 | 118.18 |
| N 714 | 8.17 | 119.70 | 8.17 | 119.49 |
| L 715 | 8.31 | 123.77 | 8.35 | 124.09 |
| Y 716 | 8.13 | 121.08 | 7.99 | 121.00 |
| Q 717 | | | | |
| T 718 | 7.86 | 109.48 | 7.87 | 109.16 |
| S 719 | | | | |
| K 720 | | | | |
| M 721 | | | | |
| K 722 | | | | |
| S 723 | | | | |

| | | | | |
|---|---|---|---|---|
| H 724 | | | | |
| G 725 | 8.35 | 110.06 | | |
| I 726 | | | | |
| F 727 | 6.37 | 111.91 | 6.42 | 111.26 |
| T 728 | 8.76 | 113.05 | 8.59 | 111.99 |
| E 729 | 7.09 | 116.58 | 7.02 | 116.75 |
| V 730 | 8.60 | 122.53 | 8.58 | 122.71 |
| N 731 | 8.84 | 124.48 | 8.86 | 124.66 |
| Q 732 | 8.65 | 121.47 | 8.65 | 121.47 |
| E 733 | 8.54 | 125.35 | 8.60 | 125.28 |
| V 734 | 8.21 | 120.62 | 8.26 | 120.61 |
| G 735 | 8.62 | 113.18 | 8.75 | 112.28 |
| E 736 | 8.23 | 115.89 | 8.26 | 116.00 |
| I 737 | 6.82 | 115.60 | 6.86 | 115.52 |
| I 738 | 8.05 | 123.90 | 8.05 | 123.62 |
| V 739 | 8.77 | 127.06 | 8.75 | 129.53 |
| A 740 | 9.13 | 132.75 | 9.18 | 133.10 |
| E 741 | 8.03 | 123.91 | 8.08 | 123.92 |
| V 742 | | | | |
| N 743 | | | | |
| H 744 | | | | |
| P 745 | | | | |
| R 746 | 8.36 | 121.75 | 8.38 | 121.64 |
| I 747 | | | | |
| A 748 | | | | |
| E 749 | | | | |
| L 750 | | | | |
| L 751 | 7.72 | 119.33 | 7.63 | 119.34 |
| T 752 | 7.33 | 119.03 | 7.32 | 118.62 |
| P 753 | | | | |
| D 754 | 7.17 | 114.92 | 7.14 | 114.90 |
| R 755 | 8.06 | 112.90 | 8.05 | 112.75 |
| V 756 | 8.11 | 123.04 | 8.11 | 123.02 |
| A 757 | 9.16 | 123.70 | 9.06 | 123.26 |
| L 758 | 8.81 | 122.60 | 8.82 | 122.89 |
| R 759 | 8.43 | 118.88 | 8.38 | 119.18 |
| K 760 | | | | |
| L 761 | 7.39 | 120.02 | 7.46 | 120.26 |
| I 762 | | | | |
| T 763 | | | | |
| K 764 | | | | |
| E 765 | | | | |
| A 766 | | | | |

**Table S6. Summary of backbone amide assignment of acetyl-CoA-bound $^{2}H$, $^{13}C$, $^{15}N$-CTD.** Chemical shifts are given in ppm. The gaps indicate missing assignments. The N-terminal polyhistidine tag is omitted. Resonances highlighted in pink indicate resonances transferred from the *apo* spectrum through the minimal shift assumption resonance translation method [19]. Among the 121 peaks picked in the $^{1}H$-$^{15}N$ HSQC spectrum, 69 were assigned to the 158 non-proline or non-N-terminal amino acids.

| Residue | $^{1}H$ | $^{15}N$ | CO | Cα | Cβ |
|---|---|---|---|---|---|
| L 601 | 8.21 | 124.55 | 174.17 | 51.38 | 38.58 |
| V 602 | 7.95 | 121.86 | 173.28 | 59.26 | 29.04 |
| Q 603 | 8.33 | 124.74 | 173.23 | 52.50 | 25.93 |
| T 604 | 8.02 | 116.29 | 171.54 | 58.80 | 66.58 |
| E 605 | 8.21 | 124.00 | 173.45 | 53.16 | 26.80 |
| V 606 | 7.98 | 125.98 | | | |
| P 607 | | | | | |
| R 608 | | | | | |
| K 609 | 8.00 | 118.84 | | 55.17 | |
| **L 610** | **6.93** | **119.26** | 174.08 | 54.67 | |
| **L 611** | **5.68** | **114.93** | 175.91 | | |
| **E 612** | **7.28** | **117.36** | 176.95 | 57.51 | 24.84 |
| E 613 | 8.27 | 122.36 | | 59.76 | |
| W 614 | | | | | |
| L 615 | | | | | |
| A 616 | | | | | |
| **M 617** | **7.63** | **120.70** | 176.25 | 55.96 | |
| W 618 | | | | | |
| S 619 | 8.17 | 116.06 | 172.10 | 55.29 | 60.91 |
| G 620 | 8.27 | 110.92 | 170.93 | 41.93 | |
| H 621 | | | | | |
| Y 622 | | | | | |
| **Q 623** | **7.35** | **115.38** | 172.23 | 53.40 | |
| L 624 | | | | | |
| **K 625** | **8.54** | **123.26** | 173.43 | 52.08 | 28.41 |
| D 626 | 7.52 | 119.94 | 172.42 | 50.56 | 39.01 |
| K 627 | 8.98 | 124.79 | 172.74 | 52.61 | 28.93 |
| L 628 | 8.11 | 123.83 | 174.39 | 52.17 | 38.28 |
| R 629 | | | | | |
| **V 630** | **9.34** | **122.29** | 172.18 | 57.52 | 29.84 |
| Q 631 | | | | | |
| L 632 | | | | | |
| R 633 | | | | | |
| P 634 | | | | | |
| Q 635 | 8.55 | 122.20 | 174.46 | 52.39 | 26.81 |
| R 636 | | | | | |
| A 637 | 8.44 | 124.07 | 176.24 | 50.76 | 14.29 |

| | | | | | |
|---|---|---|---|---|---|
| G 638 | 8.74 | 111.46 | 170.72 | 42.20 | |
| S 639 | 7.65 | 113.95 | | 54.25 | 61.72 |
| E 640 | 8.79 | 123.32 | 176.74 | | |
| V 641 | | | | | |
| L 642 | | | | | |
| E 643 | | | | | |
| L 644 | | | | | |
| G 645 | | | | | |
| I 646 | 7.93 | 121.97 | 171.03 | 51.90 | 38.60 |
| H 647 | 7.97 | 123.01 | 171.66 | 56.57 | 29.05 |
| **G 648** | **8.31** | **109.18** | | | |
| E 649 | 9.78 | 124.92 | 175.57 | 55.80 | 26.34 |
| S 650 | 8.70 | 115.31 | | 55.25 | 59.79 |
| D 651 | 8.25 | 116.92 | 173.37 | 53.46 | 36.90 |
| D 652 | 8.18 | 121.21 | 172.47 | 51.63 | 36.82 |
| K 653 | | | | | |
| L 654 | 9.15 | 126.51 | 172.03 | 51.05 | 29.52 |
| A 655 | 8.44 | 124.92 | 176.39 | 50.50 | 14.07 |
| **N 656** | **9.49** | **118.64** | 170.04 | 49.13 | 37.67 |
| V 657 | | | | | |
| I 658 | | | | | |
| **F 659** | **9.12** | **125.53** | | | |
| Q 660 | | | | | |
| P 661 | | | | | |
| I 662 | | | | | |
| Q 663 | | | | | |
| D 664 | | | | | |
| R 665 | | | | | |
| **R 666** | **7.88** | **118.92** | | 54.06 | 27.37 |
| **G 667** | **7.94** | **108.63** | 171.93 | | |
| **R 668** | **8.38** | **120.99** | 171.82 | 60.97 | |
| T 669 | | | | | |
| I 670 | | | | | |
| L 671 | | | | | |
| L 672 | | | | | |
| V 673 | | | | | |
| **R 674** | **9.05** | **128.18** | | | |
| D 675 | | | | | |
| Q 676 | | | | | |
| N 677 | | | | | |
| T 678 | | | | | |
| F 679 | | | | | |
| G 680 | 8.24 | 113.77 | 172.07 | 42.40 | |

| | | | | | |
|---|---|---|---|---|---|
| A 681 | 8.70 | 130.03 | 177.52 | 52.31 | 15.01 |
| E 682 | 9.32 | 116.22 | 173.74 | 54.90 | 25.11 |
| L 683 | | | | | |
| R 684 | | | | | |
| Q 685 | | | | | |
| K 686 | | | | | |
| R 687 | | | | | |
| L 688 | | | | | |
| **M 689** | **8.23** | **125.09** | | | |
| T 690 | | | | | |
| L 691 | | | | | |
| I 692 | | | | | |
| H 693 | | | | | |
| L 694 | | | | | |
| W 695 | | | | | |
| L 696 | | | | | |
| V 697 | | | | | |
| H 698 | | | | | |
| R 699 | | | | | |
| F 700 | | | | | |
| **K 701** | **7.44** | **118.13** | 175.87 | 56.83 | 33.34 |
| A 702 | | | | | |
| **Q 703** | **8.97** | **118.60** | | | |
| A 704 | | | | | |
| V 705 | | | | | |
| **H 706** | **9.17** | **127.56** | 170.94 | 51.90 | 29.24 |
| Y 707 | | | | | |
| V 708 | | | | | |
| T 709 | | | | | |
| P 710 | | | | | |
| T 711 | | | | | |
| **D 712** | **8.74** | **121.61** | 175.11 | 54.51 | |
| D 713 | | | | | |
| **N 714** | **8.09** | **119.88** | | 51.32 | 41.88 |
| **L 715** | **8.28** | **124.65** | | | |
| Y 716 | | | | | |
| Q 717 | | | | | |
| **T 718** | **7.99** | **107.56** | | 58.93 | 66.43 |
| S 719 | | | | | |
| K 720 | | | | | |
| M 721 | | | | | |
| K 722 | | | | | |
| S 723 | | | | | |

| | | | | | |
|---|---|---|---|---|---|
| H 724 | | | | | |
| **G 725** | **8.34** | **110.53** | 171.37 | 42.14 | |
| I 726 | | | | | |
| **F 727** | **6.33** | **111.45** | | 49.41 | |
| T 728 | | | | | |
| **E 729** | **6.95** | **117.52** | | 51.50 | 29.13 |
| V 730 | | | | | |
| **N 731** | **8.73** | **125.08** | | | |
| Q 732 | | | | | |
| E 733 | | | | | |
| V 734 | | | | | |
| G 735 | | | | | |
| **E 736** | **8.26** | **116.17** | 172.03 | 55.32 | |
| **I 737** | **6.82** | **115.36** | | | |
| **I 738** | **8.14** | **124.62** | 174.40 | 56.97 | |
| V 739 | | | | | |
| A 740 | | | | | |
| **E 741** | **7.99** | **124.16** | | | |
| V 742 | | | | | |
| N 743 | | | | | |
| H 744 | | | | | |
| P 745 | | | | | |
| **R 746** | **8.34** | **122.28** | 174.40 | 53.20 | 27.08 |
| I 747 | | | | | |
| A 748 | | | | | |
| E 749 | | | | | |
| L 750 | | | | | |
| **L 751** | **7.63** | **119.64** | 181.18 | 51.25 | 37.34 |
| T 752 | 7.27 | 119.08 | 173.22 | 59.75 | 65.76 |
| P 753 | | | | | |
| D 754 | 7.18 | 115.41 | 173.97 | 51.95 | 36.63 |
| R 755 | 7.98 | 113.23 | 173.28 | 55.26 | |
| V 756 | 8.02 | 123.57 | 176.07 | 64.32 | 27.45 |
| A 757 | 8.98 | 123.72 | 178.54 | 51.52 | |
| L 758 | 8.68 | 122.76 | 175.57 | 54.47 | |
| R 759 | 8.36 | 119.67 | | | |
| K 760 | | | | | |
| **L 761** | **7.39** | **120.69** | 176.57 | 54.35 | 36.61 |
| I 762 | | | | | |
| T 763 | | | | | |
| K 764 | | | | | |
| E 765 | | | | | |
| A 766 | 7.72 | 129.80 | 179.38 | 50.37 | 16.71 |

**Table S7. Transfer of $^1$H and $^{15}$N assignments between *apo* $^2$H, $^{13}$C, $^{15}$N-CTD and acetyl-CoA bound $^2$H, $^{13}$C, $^{15}$N-CTD in TROSY HSQC.** Chemical shifts are given in ppm. Gaps indicate missing assignments or transfers. Resonances highlighted in pink indicate resonances transferred from the other spectrum through the minimal shift assumption resonance translation method [19].

| | ***Apo* $^2$H, $^{15}$N-CTD** | | **Acetyl-CoA-bound $^2$H, $^{15}$N-CTD** | |
|---|---|---|---|---|
| **Residue** | **$^1$H** | **$^{15}$N** | **$^1$H** | **$^{15}$N** |
| L 601 | 8.15 | 123.54 | 8.21 | 124.55 |
| V 602 | 7.93 | 121.66 | 7.95 | 121.86 |
| Q 603 | 8.32 | 124.70 | 8.33 | 124.74 |
| T 604 | 8.01 | 116.15 | 8.02 | 116.29 |
| E 605 | 8.22 | 124.05 | 8.21 | 124.00 |
| V 606 | 8.09 | 124.84 | 7.98 | 125.98 |
| P 607 | | | | |
| R 608 | | | | |
| K 609 | 8.09 | 118.04 | 8.00 | 118.84 |
| L 610 | 6.99 | 119.32 | **6.93** | **119.26** |
| L 611 | 5.69 | 114.93 | **5.68** | **114.93** |
| E 612 | 7.23 | 117.30 | **7.28** | **117.36** |
| E 613 | | | 8.27 | 122.36 |
| W 614 | | | | |
| L 615 | | | | |
| A 616 | | | | |
| M 617 | 7.72 | 121.04 | **7.63** | **120.70** |
| W 618 | | | | |
| S 619 | | | 8.17 | 116.06 |
| G 620 | **8.24** | **110.87** | 8.27 | 110.92 |
| H 621 | | | | |
| Y 622 | | | | |
| Q 623 | 7.43 | 115.84 | **7.35** | **115.38** |
| L 624 | 8.04 | 119.59 | | |
| K 625 | 8.51 | 122.21 | **8.54** | **123.26** |
| D 626 | 7.54 | 119.63 | 7.52 | 119.94 |
| K 627 | 8.90 | 124.86 | 8.98 | 124.79 |
| L 628 | | | 8.11 | 123.83 |
| R 629 | | | | |
| V 630 | 9.32 | 122.27 | **9.34** | **122.29** |
| Q 631 | | | | |
| L 632 | | | | |
| R 633 | | | | |
| P 634 | | | | |
| Q 635 | | | 8.55 | 122.20 |
| R 636 | 7.44 | 111.69 | | |
| A 637 | 8.44 | 124.57 | 8.44 | 124.07 |
| G 638 | 8.76 | 111.59 | 8.74 | 111.46 |

| | | | | |
|---|---|---|---|---|
| S 639 | 7.73 | 114.20 | 7.65 | 113.95 |
| E 640 | 8.73 | 124.04 | 8.79 | 123.32 |
| V 641 | | | | |
| L 642 | 8.55 | 128.10 | | |
| E 643 | 8.68 | 118.61 | | |
| L 644 | | | | |
| G 645 | | | | |
| I 646 | **7.91** | **121.94** | 7.93 | 121.97 |
| H 647 | **7.96** | **123.00** | 7.97 | 123.01 |
| G 648 | 8.26 | 108.74 | **8.31** | **109.18** |
| E 649 | 9.65 | 124.62 | 9.78 | 124.92 |
| S 650 | 8.63 | 114.98 | 8.70 | 115.31 |
| D 651 | 8.25 | 116.75 | 8.25 | 116.92 |
| D 652 | 8.22 | 121.43 | 8.18 | 121.21 |
| K 653 | | | | |
| L 654 | **9.20** | **126.91** | 9.15 | 126.51 |
| A 655 | | | 8.44 | 124.92 |
| N 656 | 9.31 | 118.45 | **9.49** | **118.64** |
| V 657 | | | | |
| I 658 | | | | |
| F 659 | 9.05 | 125.66 | **9.12** | **125.53** |
| Q 660 | | | | |
| P 661 | | | | |
| I 662 | | | | |
| Q 663 | 7.95 | 119.45 | | |
| D 664 | 8.62 | 123.23 | | |
| R 665 | 8.29 | 117.05 | | |
| R 666 | 7.88 | 118.52 | **7.88** | **118.92** |
| G 667 | 8.02 | 109.00 | **7.94** | **108.63** |
| R 668 | 8.43 | 121.76 | **8.38** | **120.99** |
| T 669 | 8.83 | 120.01 | | |
| I 670 | | | | |
| L 671 | | | | |
| L 672 | | | | |
| V 673 | | | | |
| R 674 | 9.04 | 129.07 | **9.05** | **128.18** |
| D 675 | | | | |
| Q 676 | | | | |
| N 677 | | | | |
| T 678 | | | | |
| F 679 | | | | |
| G 680 | **8.22** | **113.71** | 8.24 | 113.77 |
| A 681 | **8.75** | **131.03** | 8.70 | 130.03 |

| | | | | |
|---|---|---|---|---|
| E 682 | **9.23** | **116.52** | 9.32 | 116.22 |
| L 683 | | | | |
| R 684 | | | | |
| Q 685 | | | | |
| K 686 | | | | |
| R 687 | | | | |
| L 688 | | | | |
| M 689 | 8.21 | 124.95 | **8.23** | **125.09** |
| T 690 | | | | |
| L 691 | | | | |
| I 692 | | | | |
| H 693 | | | | |
| L 694 | | | | |
| W 695 | | | | |
| L 696 | | | | |
| V 697 | | | | |
| H 698 | | | | |
| R 699 | | | | |
| F 700 | | | | |
| K 701 | 7.36 | 119.22 | **7.44** | **118.13** |
| A 702 | 7.67 | 118.46 | | |
| Q 703 | 9.14 | 117.69 | **8.97** | **118.60** |
| A 704 | 7.59 | 117.49 | | |
| V 705 | 9.24 | 120.99 | | |
| H 706 | 9.28 | 127.34 | **9.17** | **127.56** |
| Y 707 | 9.48 | 122.68 | | |
| V 708 | 7.31 | 118.15 | | |
| T 709 | | | | |
| P 710 | | | | |
| T 711 | 7.70 | 113.31 | | |
| D 712 | 8.82 | 121.68 | **8.74** | **121.61** |
| D 713 | 8.64 | 118.85 | | |
| N 714 | 8.08 | 120.15 | **8.09** | **119.88** |
| L 715 | 8.28 | 124.42 | **8.28** | **124.65** |
| Y 716 | 8.04 | 121.53 | | |
| Q 717 | | | | |
| T 718 | 7.98 | 108.47 | **7.99** | **107.56** |
| S 719 | | | | |
| K 720 | | | | |
| M 721 | | | | |
| K 722 | | | | |
| S 723 | | | | |
| H 724 | | | | |

| | | | | |
|---|---|---|---|---|
| G 725 | 8.33 | 110.52 | **8.34** | **110.53** |
| I 726 | | | | |
| F 727 | 6.33 | 112.63 | **6.33** | **111.45** |
| T 728 | 8.71 | 113.72 | | |
| E 729 | 7.03 | 117.15 | **6.95** | **117.52** |
| V 730 | 8.55 | 123.03 | | |
| N 731 | 8.78 | 124.95 | **8.73** | **125.08** |
| Q 732 | 8.60 | 122.16 | | |
| E 733 | | | | |
| V 734 | | | | |
| G 735 | 8.56 | 113.77 | | |
| E 736 | 8.18 | 116.53 | **8.26** | **116.17** |
| I 737 | 6.81 | 116.05 | **6.82** | **115.36** |
| I 738 | 8.17 | 124.31 | **8.14** | **124.62** |
| V 739 | 8.69 | 127.65 | | |
| A 740 | 9.04 | 133.26 | | |
| E 741 | 7.98 | 124.26 | **7.99** | **124.16** |
| V 742 | | | | |
| N 743 | | | | |
| H 744 | | | | |
| P 745 | | | | |
| R 746 | 8.32 | 122.21 | **8.34** | **122.28** |
| I 747 | | | | |
| A 748 | | | | |
| E 749 | | | | |
| L 750 | | | | |
| L 751 | 7.66 | 119.88 | **7.63** | **119.64** |
| T 752 | 7.27 | 119.49 | 7.27 | 119.08 |
| P 753 | | | | |
| D 754 | 7.11 | 115.41 | 7.18 | 115.41 |
| R 755 | 8.01 | 113.42 | 7.98 | 113.23 |
| V 756 | 8.06 | 123.55 | 8.02 | 123.57 |
| A 757 | 9.08 | 124.09 | 8.98 | 123.72 |
| L 758 | 8.73 | 123.07 | 8.68 | 122.76 |
| R 759 | 8.38 | 119.40 | 8.36 | 119.67 |
| K 760 | | | | |
| L 761 | 7.33 | 120.45 | **7.39** | **120.69** |
| I 762 | | | | |
| T 763 | | | | |
| K 764 | | | | |
| E 765 | | | | |
| A 766 | **7.76** | **129.71** | 7.72 | 129.80 |

**Table S8. Interactions between unique helical substructure (UHS) residues and other monomeric chains in *apo* ICL2.** Interacting residues were taken from the PISA analysis [14] of *apo* ICL2 (PDB: 6EDW [17]). The residues are listed as "chain: residue name and number [interacting atom]". Interactions absent in the acetyl-CoA-bound form are highlighted in orange, interactions between UHS and the C-terminal domain (CTD) residues in magenta, and interactions between UHS and the flexible linker in blue. The interactions of UHS with the CTS and linker are also absent in the acetyl-CoA-bound ICL2.

| Hydrogen Bonds | | | Salt Bridges | | |
|---|---|---|---|---|---|
| Residue 1 | Distance (Å) | Residue 2 | Residue 1 | Distance (Å) | Residue 2 |
| D:ARG 584 [NH2] | 3.81 | B:GLU 302 [OE1] | D:ARG 584 [NH2] | 3.81 | B:GLU 302 [OE1] |
| D:ARG 584 [NH1] | 2.63 | B:ASP 348 [OD1] | D:ARG 584 [NH2] | 3.22 | B:ASP 348 [OD1] |
| **D:LYS 589 [NZ]** | **3.54** | **B:ASP 352 [O ]** | D:ARG 584 [NH1] | 2.63 | B:ASP 348 [OD1] |
| **D:LYS 589 [NZ]** | **2.63** | **B:ASP 352 [OD1]** | **D:LYS 589 [NZ]** | **2.63** | **B:ASP 352 [OD1]** |
| **D:LYS 589 [NZ]** | **3.31** | **B:GLU 355 [OE1]** | **D:LYS 589 [NZ]** | **3.31** | **B:GLU 355 [OE1]** |
| **D:GLY 592 [N ]** | **2.75** | **B:GLU 363 [OE2]** | **D:ARG 699 [NH1]** | **3.37** | **B:GLU 385 [OE1]** |
| **D:ARG 608 [NH2]** | **2.58** | **B:GLU 385 [O ]** | **D:ARG 699 [NH2]** | **3.75** | **B:GLU 387 [OE2]** |
| **D:ARG 608 [N ]** | **2.82** | **B:GLU 385 [OE1]** | D:GLU 302 [OE1] | 3.80 | B:ARG 584 [NH2] |
| **D:ARG 699 [NH1]** | **3.37** | **B:GLU 385 [OE1]** | D:ASP 348 [OD1] | 3.64 | B:ARG 584 [NH2] |
| **D:ARG 699 [NH2]** | **3.75** | **B:GLU 387 [OE2]** | D:ASP 348 [OD1] | 3.01 | B:ARG 584 [NH1] |
| D:GLU 302 [OE1] | 3.80 | B:ARG 584 [NH2] | D:ASP 348 [OD2] | 3.82 | B:ARG 584 [NH2] |
| D:ASP 348 [OD1] | 3.01 | B:ARG 584 [NH1] | **D:ASP 352 [OD1]** | **2.83** | **B:LYS 589 [NZ]** |
| **D:ASP 352 [OD1]** | **2.83** | **B:LYS 589 [NZ]** | **D:ASP 352 [OD2]** | **3.97** | **B:LYS 589 [NZ]** |
| **D:SER 356 [OG]** | **2.94** | **B:LYS 589 [NZ]** | **D:ASP 377 [OD1]** | **2.81** | **B:HIS 600 [ND1]** |
| **D:GLU 363 [OE2]** | **2.73** | **B:GLY 592 [N ]** | **D:ASP 377 [OD2]** | **3.94** | **B:HIS 600 [ND1]** |
| **D:ASP 377 [OD1]** | **2.81** | **B:HIS 600 [ND1]** | **D:GLU 385 [OE1]** | **3.92** | **B:ARG 608 [NE]** |
| **D:ASP 377 [OD2]** | **2.89** | **B:LEU 601 [N ]** | C:LYS 285 [NZ] | 2.75 | B:GLU 267 [OE1] |
| **D:ASP 377 [OD2]** | **2.85** | **B:THR 604 [OG1]** | C:LYS 285 [NZ] | 3.07 | B:GLU 267 [OE2] |
| **D:GLU 385 [O ]** | **2.84** | **B:ARG 608 [NH2]** | **C:HIS 406 [ND1]** | **3.75** | **B:GLU 315 [OE1]** |
| **D:GLU 385 [OE1]** | **2.97** | **B:ARG 608 [N ]** | **C:HIS 406 [ND1]** | **3.96** | **B:GLU 315 [OE2]** |
| C:ASN 304 [ND2] | 2.97 | B:ARG 39 [O ] | C:GLU 267 [OE1] | 2.83 | B:LYS 285 [NZ] |
| C:TYR 309 [OH] | 2.68 | B:ASP 227 [OD2] | C:GLU 267 [OE2] | 3.04 | B:LYS 285 [NZ] |
| C:LYS 285 [NZ] | 2.95 | B:ALA 265 [O ] | C:GLU 315 [OE1] | 3.96 | B:HIS 406 [ND1] |
| C:LYS 285 [NZ] | 2.75 | B:GLU 267 [OE1] | C:GLU 315 [OE2] | 3.75 | B:HIS 406 [ND1] |
| C:ARG 268 [NH1] | 2.95 | B:LEU 308 [O ] | | | |
| C:ARG 268 [NH2] | 2.77 | B:LEU 308 [O ] | | | |
| C:ARG 39 [O ] | 2.88 | B:ASN 304 [ND2] | | | |
| C:ASP 227 [OD2] | 2.67 | B:TYR 309 [OH] | | | |
| C:ALA 265 [O ] | 2.93 | B:LYS 285 [NZ] | | | |
| C:GLU 267 [OE1] | 2.83 | B:LYS 285 [NZ] | | | |
| C:LEU 308 [O ] | 3.08 | B:ARG 268 [NH1] | | | |
| C:LEU 308 [O ] | 2.88 | B:ARG 268 [NH2] | | | |
| C:GLU 315 [OE1] | 2.76 | B:SER 404 [OG] | | | |

**Table S9. Interactions between unique helical substructure residues and other monomeric chains in acetyl-CoA-bound ICL2.** Interacting residues were taken from the PISA analysis [14] of acetyl-CoA-bound ICL2 (PDB: 6EE1 [17]). The residues are listed as "chain: residue name and number [interacting atom]". Interactions absent in the *apo* ICL2 form are highlighted in orange.

| Hydrogen Bonds | | | Salt Bridges | | |
|---|---|---|---|---|---|
| Residue 1 | Distance (Å) | Residue 2 | Residue 1 | Distance (Å) | Residue 2 |
| D:ARG 584 [NH2] | 3.79 | B:ASP 348 [OD2] | D:ARG 584 [NH1] | 3.27 | B:ASP 348 [OD1] |
| D:GLU 302 [OE2] | 3.06 | B:ARG 584 [NH2] | D:ARG 584 [NH2] | 3.79 | B:ASP 348 [OD2] |
| C:ASN 304 [ND2] | 2.84 | B:ARG 39 [O ] | D:ARG 584 [NH1] | 3.55 | B:ASP 348 [OD2] |
| C:TYR 309 [OH] | 2.84 | B:ASP 227 [OD2] | D:GLU 302 [OE2] | 3.06 | B:ARG 584 [NH2] |
| C:LYS 285 [NZ] | 2.95 | B:ALA 265 [O ] | D:ASP 348 [OD1] | 3.51 | B:ARG 584 [NH1] |
| C:LYS 285 [NZ] | 2.78 | B:GLU 267 [OE1] | D:ASP 348 [OD2] | 3.79 | B:ARG 584 [NH1] |
| C:ARG 268 [NH2] | 2.69 | B:LEU 308 [O ] | C:LYS 285 [NZ] | 2.78 | B:GLU 267 [OE1] |
| C:SER 404 [OG] | 3.59 | B:GLU 315 [OE1] | C:LYS 285 [NZ] | 3.07 | B:GLU 267 [OE2] |
| **C:ARG 44 [NH1]** | **3.63** | **B:GLN 345 [OE1]** | **C:ARG 264 [NH2]** | **3.37** | **B:GLU 424 [OE1]** |
| C:ARG 39 [O ] | 2.89 | B:ASN 304 [ND2] | **C:ARG 264 [NH2]** | **3.09** | **B:GLU 424 [OE2]** |
| C:ASP 227 [OD2] | 2.78 | B:TYR 309 [OH] | C:GLU 267 [OE1] | 2.88 | B:LYS 285 [NZ] |
| C:ALA 265 [O ] | 2.86 | B:LYS 285 [NZ] | C:GLU 267 [OE2] | 2.94 | B:LYS 285 [NZ] |
| C:GLU 267 [OE1] | 2.88 | B:LYS 285 [NZ] | **C:GLU 424 [OE1]** | **3.93** | **B:ARG 264 [NE]** |
| C:GLU 267 [OE2] | 2.94 | B:LYS 285 [NZ] | **C:GLU 424 [OE1]** | **3.63** | **B:ARG 264 [NH2]** |
| C:LEU 308 [O ] | 2.81 | B:ARG 268 [NH2] | **C:GLU 424 [OE2]** | **3.99** | **B:ARG 264 [NE]** |
| C:GLU 315 [OE1] | 3.12 | B:SER 404 [OG] | **C:GLU 424 [OE2]** | **3.26** | **B:ARG 264 [NH2]** |

**Table S10. Timescale for molecular dynamics simulations.**

| Sim # | *apo* dimer | *apo* monomer | CoA dimer | CoA monomer | CoA remove monomer |
|---|---|---|---|---|---|
| 1 | 326.3 ns | 275.9 ns | 307.4 ns | 368 ns | 423.7 ns |
| 2 | 325.9 ns | 275.7 ns | 307.1 ns | 367.2 ns | 367.3 ns |
| 3 | 323.8 ns | 276.0 ns | 306.8 ns | 367.1 ns | 460.7 ns |
